# Spatial transcriptomic analysis of muscle biopsy from treatment-naive juvenile dermatomyositis patients reveals mitochondrial abnormalities despite disease-related interferon driven signature

**DOI:** 10.1101/2025.06.03.657601

**Authors:** Aris E Syntakas, Melissa Kartawinata, Nia M L Evans, Huong D Nguyen, Charalampia Papadopoulou, Muthana Al Obaidi, Clarissa Pilkington, Yvonne Glackin, Christopher B Mahoney, Adam P Croft, Simon Eaton, Mario Cortina-Borja, Olumide Ogunbiyi, Ashirwad Merve, Lucy R Wedderburn, Meredyth G Ll Wilkinson, the JDCBS

## Abstract

**Objectives:** This study aimed to investigate the spatial transcriptomic landscape of muscle tissue from treatment-naïve juvenile dermatomyositis (JDM) patients in comparison to healthy paediatric muscle tissue.

**Methods:** Muscle biopsies from three JDM patients and three age-matched controls were analysed using the Nanostring GeoMx® Digital Spatial Profiler. Regions of interest were selected based on muscle fibres without immune cells, immune cell infiltration and CD68+ macrophage enrichment. Differential gene expression, pathway analysis and pathways clustering analysis were conducted. Key findings were validated in 19 cases of JDM using immunohistochemistry and chemical stains, and a bulk RNAseq dataset of four cases of JDM.

**Results:** JDM muscle tissues exhibited significant interferon pathway activation and mitochondrial dysfunction compared to controls. A 15-gene interferon signature was significantly elevated in JDM muscle and macrophage-enriched regions, correlating with clinical weakness. In contrast, mitochondrial dysregulation, characterized by downregulated respiratory chain pathways, was present regardless of interferon activity or muscle strength. The interferon-driven and mitochondrial signatures were replicated in an independent RNAseq dataset from JDM muscle; lack of association between interferon signature and mitochondrial dysregulation was validated in 19 cases by conventional staining methods. Clustering analysis revealed distinct transcriptomic profiles between JDM and control tissues, as well as between JDM patients with varying clinical phenotypes.

**Conclusions:** This study highlights mitochondrial dysfunction as a consistent pathological feature in JDM muscle, which may be independent of interferon-driven inflammation. These findings highlight the potential for mitochondrial-targeted therapies in JDM management and emphasise the need for further studies to explore their therapeutic value.

**KEY MESSAGES:** What is already known on this topic

- Juvenile dermatomyositis (JDM) involves interferon-driven inflammation and immune-mediated muscle damage.
- Mitochondrial abnormalities in blood immune cells persist despite treatment and contribute to disease pathology.

What this study adds

- Mitochondrial dysfunction is present in both muscle fibres and tissue-infiltrating immune cells within JDM muscle.
- These abnormalities are detectable even in clinically less severe muscle weakness.
- Degree of mitochondrial abnormality at transcript and protein level may be independent of strength of IFN-driven signal.
- Mitochondrial dysregulation detected at transcriptional level correlates with abnormal transcription of muscle (sarcomere) and the subcellular peroxisome organelle.

How this study might affect research, practice, or policy

- Targeting mitochondrial dysfunction could enhance treatment outcomes for JDM, especially in patients whose disease is refractory to current therapies.
- Insights from this study support the development of stratification tools to detect aspects of pathology which are not tightly correlated with IFN-driven pathology.

## INTRODUCTION

Idiopathic inflammatory myopathies of childhood, of which the most common is juvenile dermatomyositis (JDM), are severe paediatric autoimmune conditions characterised by chronic inflammation of muscle, skin and in some cases major organs. A clear understanding of pathogenic mechanisms at the sites of tissue damage is lacking, and there is a significant unmet need for more targeted treatments^1, 2^. We have previously detailed transcriptional changes detectable in blood immune cells in JDM^3^. We identified that blood CD14+ monocytes from patients with active disease have severe mitochondrial dysfunction. Further we showed that alterations in mitochondrial biology and morphology in monocytes led to accumulation of oxidized mitochondrial DNA (oxmtDNA), which was able to trigger IFN-signalling pathways. Interestingly, while the well-characterised IFN-driven signature detectable at diagnosis was reduced on treatment, we observed that mitochondrial abnormalities detected in peripheral blood monocytes persisted despite treatment.

Our previous studies on JDM muscle tissue obtained prior to treatment showed that pro-inflammatory, CD68+ tissue macrophages are frequently detectable^4,5^ and that CD68+ cell infiltration correlates with weakness^6^. Macrophage infiltration may be throughout the endomysium, or in peri-vascular clusters, where they are frequently co-localized with infiltrating T cells^5,6^. Abnormal macrophage polarisation has been implicated in perpetuating inflammation and tissue damage in myopathies^7–9^. A recent study in a myositis animal model suggests a self-perpetuating loop between mitochondrial dysfunction and inflammation^10^.

To start to define at tissue level the complex interactions between immune and muscle cells in JDM-affected muscle tissue, we employed spatial transcriptomics. This method enables in-situ gene expression analysis while preserving critical tissue architecture, allowing precise localization of cell types and their potential involvement in the pathogenic processes associated with myositis^11^. We employed the Nanostring GeoMx® Digital Spatial Profiler (DSP) to interrogate the transcriptome of muscle tissue sections from JDM patients and age-matched healthy controls. GeoMx facilitates the staining, selection and sequencing of specific spatial niches within the tissue known as Regions of Interest (ROIs)^12^. We opted to focus initially on CD68+-enriched areas in the analysis of inflammatory infiltrate.

We confirm that JDM muscle tissue exhibits significant upregulation of interferon (IFN) pathways and dysregulation of mitochondrial pathways, particularly affecting the electron transport chain. By comparing tissue areas, we show that muscle weakness correlates with increased IFN signalling in both muscle- and CD68+-cell-derived transcriptome. Importantly mitochondrial abnormalities were clearly demonstrated even in muscle with only moderate clinical weakness, and these were detected in both muscle fibres as well as in the CD68+ infiltrate. We replicated our findings in another cohort, analysing bulk RNAseq data, and validated the lack of correlation between IFN-driven changes and mitochondrial abnormality at protein level in a larger cohort. We believe that our study is the first to provide spatial transcriptional insights into JDM pathogenesis, and will facilitate significant steps towards the delineation of the contributions of immune and muscle cells, to JDM pathology. Our results suggest that mitochondrial abnormalities, that we previously defined in blood cells, are also present in both muscle and infiltrating monocyte/macrophages in JDM even before significant clinical weakness is apparent.

## METHODS

all methods are provided in supplementary information

## RESULTS

### Patients and controls

Clinical and demographic features of the 3 JDM patients and controls analysed by transcriptome profiling are shown in Table 1. Of note, two of the patients were significantly weak at time of biopsy as assessed by the CMAS and MMT8 while the third was less weak but had more severe skin disease activity. Two JDM patients tested positive for TIF1γ, while the third patient was negative for MSA. For histological analysis, a total of 19 cases were included: the 3 transcriptome-profiled patients, along with an additional 16 JDM cases (Supplementary, Table S1).

**Table 1.** Demographics and clinical characteristics of JDM patients and controls included in the study.

| Characteristic | Value |  |  |  |  |  |
| --- | --- | --- | --- | --- | --- | --- |
| Sample ID | JDM1 | JDM2 | JDM3 | HC1 | HC2 | HC3 |
| Age at Sample, years | 5.83 | 3.58 | 4.75 | 6.28 | 10.02 | 4.81 |
| Sex | F | F | F | M | M | F |
| Disease duration at time of biopsy, months | 1.05 | 5.91 | 3.28 | NA | NA | NA |
| Myositis specific antibody | TIF1-γ | TIF1-γ | Negative | NA | NA | NA |
| Disease activity at time of biopsy: |  |  |  |  |  |  |
| Physician global assessment score, PGA (0.0-10.0 cm) | 8.0 | 7.8 | 6.3 | NA | NA | NA |
| CMAS (0-52) | 30 | 12 | 52 | NA | NA | NA |
| MMT8 (0-80) | 44 | 42 | 78 | NA | NA | NA |
| CK, U/L | 2611 | 125 | 131 | NA | NA | NA |
| sDAS* | 4 | 3 | 5 | NA | NA | NA |
| Muscle biopsy score data** |  |  |  |  |  |  |
| Inflammatory domain (0-12) | 7 | 8 | 7 | NA | NA | NA |
| Muscle fibre domain (0-10) | 8 | 7 | 7 | NA | NA | NA |
| Vascular domain (0-3) | 2 | 1 | 1 | NA | NA | NA |
| Connective tissue domain (0-2) | 1 | 2 | 1 | NA | NA | NA |
| Total Biopsy Score (0-27) | 18 | 18 | 16 | NA | NA | NA |
| Histopathology visual analogue score (0.0-10.0) | 7.0 | 8.0 | 7.0 | NA | NA | NA |

\*Skin Disease activity score, or modified skin DAS as defined in Lam et al. (2011)<sup>13</sup>
\*\*Data generated using validated JDM muscle biopsy score tool from Varsani et al. (2015)<sup>6</sup>

### Selection of regions of interest (ROIs) for analysis of areas with or without leukocyte infiltration in JDM muscle biopsies

To determine the dominant pathways which are altered in the muscle of patients with JDM compared to healthy muscle and to test whether mitochondrial pathways are altered in muscle tissue early in JDM, we used spatial transcriptome analysis. Imaging of quadriceps muscle biopsy sections from all 6 cases was performed on the Nanostring GeoMx® scanning platform. Immunofluorescence staining was performed to enable selection of ROIs for sequencing using a nuclear marker and morphology markers laminin, CD45, and CD68, for identification of muscle fibres, leukocytes, and macrophages respectively. For focused transcriptome analysis of infiltrating macrophages, segmentation was used, as described^12^. Figure 1A shows control muscle tissue, where no significant immune cell infiltration or inflammation is observed; Figure 1B shows JDM muscle ROI without immune cells, while Figure 1C reveals an area with substantial inflammatory infiltration (CD45+), which was not observed in the control samples. Figure 1D shows an area with infiltrating CD68+ cells within JDM muscle with Figure 1E further illustrating the ‘segmented’ view of CD68+ cells, allowing for transcriptomic profiling of this specific cell type. Thus, we selected three different ROI types for sequencing analysis: muscle fibre, muscle with immune cell infiltration, and CD68-enriched, generating a total of 28 ROIs. The breakdown of the ROIs selected is shown in Table S2.

**Figure 1.**
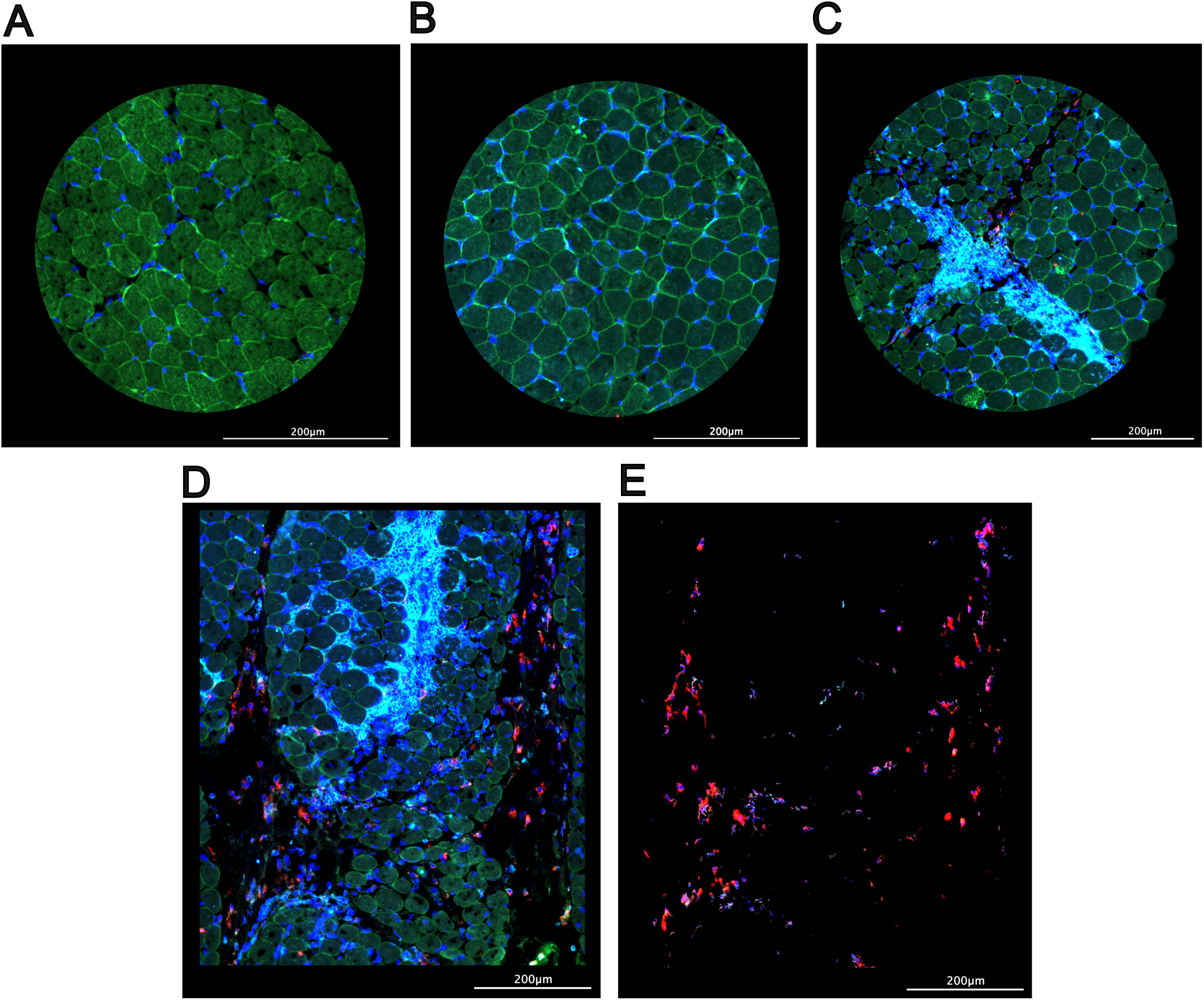
Immunofluorescent imaging of the muscle biopsy regions of interest (ROIs). (A-E) Representative fluorescent images of ROI types selected for sequence analysis and associated staining markers: (A) Control muscle; (B) JDM muscle ROI with no infiltrating immune cells; (C) JDM muscle enriched with immune cells (CD45+); (D) CD68+ macrophage-enriched region. (E) Segmented CD68+ cells in the same region shown in D. Staining antibodies: laminin (green), DNA nuclear stain (blue), CD68 (red), and CD45 (cyan), indicating muscle fibres, nuclei, macrophages, and leukocytes, respectively. B-E are derived from patient JDM3.

Histological analysis of the muscle biopsy sections is presented in Figure S1. All 3 control cases were found to have no significant pathology or inflammation, with no infiltration by CD3+ or CD68+ immune cells, confirmed by a senior histopathologist (AM)^6, 14^. None of the 3 control cases had a genetic abnormality in known mitochondrial-related diseases. A representative H&E-stained section from a control case is shown in Figure S1A. Active inflammation in a representative JDM case is demonstrated by H&E staining (Figure S1B), CD3 (Figure S1C), CD68 (Figure S1D), and CD20 staining (Figure S1E). These classical signs of JDM muscle inflammation are reflected in the biopsy scores (inflammatory domain)^6^, Table 1.

### Principal component analysis confirms distinct transcriptomic differences between ROI types and JDM vs. control tissues

Initial data quality checks were performed on the Nanostring platform, assessing tissue and sequencing quality, as well as the performance of negative control probes. None of the 28 ROIs or control probes were flagged as unsuitable for analysis. Further quality control and normalization techniques were applied to address technical variation across data generated from ROIs. Relative log expression (RLE) plots were used to visualize effects of normalization: Figure S2A shows unnormalized data, with Figure S2B displaying log counts per million (logCPM) normalized data. This normalization method, integrated into the voom function of the limma-voom pipeline^15^, effectively centred the median expression levels of all ROIs around zero. This indicates successful removal of unwanted technical variation, enhancing comparability across samples.

We employed principal component analysis (PCA) to assess variance within the dataset, between patients, and between types of ROIs (Figure S2C-D). Initial PCA analysis by patient revealed that all data from JDM patients (22 ROIs) clustered into two clusters, with clear separation of JDM patients from control samples (Figure S2C). When data were labelled by ROI type, the two distinct clusters of data from JDM biopsies were clarified by cell type. As expected CD68+-selected ROIs (JDM samples only) formed a separate cluster (Figure S2D blue symbols). Interestingly, JDM muscle only (Figure S2D red symbols) and JDM muscle plus-immune ROIs (Figure S2D yellow symbols) clustered together (Figure S2D). Note that segmentation was not performed on CD45+ cells. Again, control muscle ROIs (Figure S2D pink symbols) clustered away from all JDM data.

### Differential gene expression analysis confirms a high interferon-driven signature and demonstrates abnormal mitochondrial gene signature in JDM muscle compared to control muscle

To investigate differences in the gene expression profiles of JDM muscle compared to control muscle, the data from muscle-only ROIs were initially used in the limma-voom differential gene expression (DGE) analysis pipeline. The analysis revealed 448 genes which were significantly differentially expressed in JDM muscle (|Log2FC| ≥ 0.58 and adjusted p-value ≤ 0.05), of which 336 genes were identified as upregulated and 112 as downregulated in JDM muscle compared to control samples (Figure 2A). Pathway enrichment using Over-Representation Analysis (ORA) on significant differentially expressed genes (DEG) was used to identify dysregulated pathways in JDM muscle. The GO Cellular Component (CC) gene sets were initially used to identify differentially expressed pathways. The GO CC annotations detail the subcellular structures and complexes involved, making this appropriate for highlighting mitochondrial functional pathways. Conversely, the GO Biological Process (BP) gene sets were then used to capture dynamic biological activities and signalling processes, such as immune response mechanisms, which are central to JDM. The first ORA (GO CC) indicated 101 significantly differentially expressed pathways in JDM muscle compared to controls (adjusted p value ≤ 0.05) while the second, (GO BP), gave 310 significant pathways.

**Figure 2.**
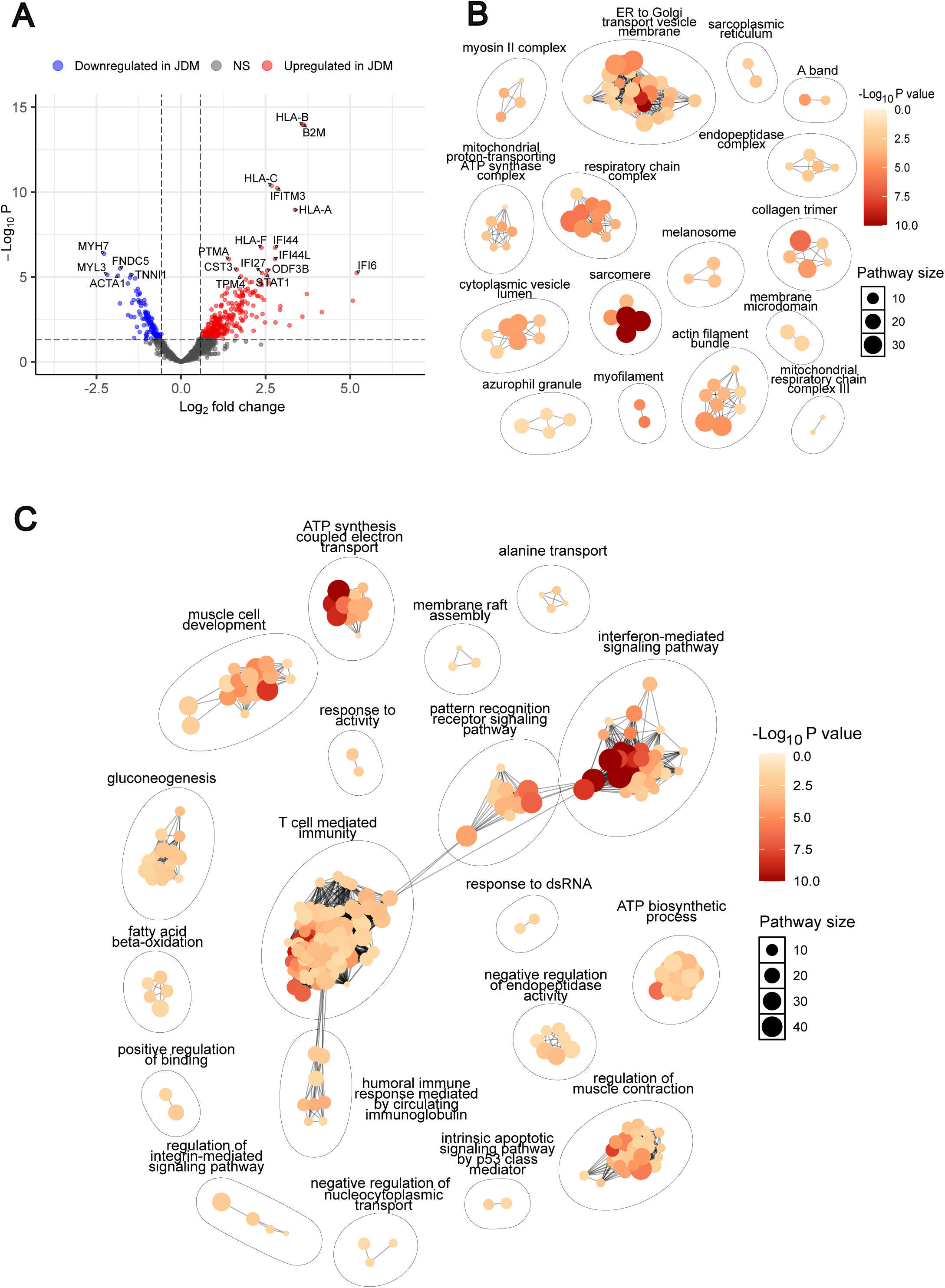
Differential gene expression analysis demonstrates high interferon-driven signature and abnormal mitochondrial gene signature in JDM compared to control muscle. (A) Volcano plot illustrating differentially expressed genes (DEGs) between JDM and control muscle ROIs. Genes with an adjusted p-value ≤ 0.05 and |log2FC| ≥ 0.58 are considered statistically significant (n = 448). Red points represent significantly upregulated genes, blue significantly downregulated genes. The 20 genes with the smallest adjusted p-values were annotated. (B) Network cluster plot of enriched Gene Ontology Cellular Component (CC) pathways among the 448 DEGs, generated using the aPEAR R package. Each cluster represents groups of similar cellular components, derived from Gene Ontology terms. Pathways with an adjusted p-value ≤ 0.05 were included in the clustering and network analysis. (C) Network cluster plot of enriched Gene Ontology Biological Process (BP) pathways among the 448 DEGs, generated using the aPEAR R package. Each cluster represents groups of related biological processes, highlighting functional pathways involved in JDM. Pathways with an adjusted p-value ≤ 0.05 were included in this clustering and network analysis. For (B) and (C), larger dots represent pathways associated with a higher number of significant genes (shown in key, pathway size). Dot colour intensity corresponds to significance, with deeper red shades indicating smaller p-values (shown in colour bar, adjusted p value). NS = not significant; FC = fold change.

To simplify interpretation of these numerous, often overlapping pathways, aPEAR cluster network analysis was performed^16^, resulting in the construction of plots that visually organize and group related pathways into coherent clusters, for each of the two GO gene sets that were used. The GO CC network highlighted several clusters of mitochondria-related pathways, including clusters such as the “respiratory chain complex”, the “mitochondrial proton-transporting ATP synthase complex”, and the “mitochondrial respiratory chain complex III” (Figure 2B). Interestingly, pathways related to ER to Golgi protein transport were also differentially expressed in JDM muscle (Figure 2B). The GO BP network highlighted clusters of interferon and immune related pathways, with the most prominent clusters being “interferon-mediated signalling pathway”, “pattern recognition receptor signalling pathway”, and “T cell mediated immunity”. In addition, several clusters related to muscle development and function were noted, including “muscle cell development” and “regulation of muscle contraction” (Figure 2C).

We were intrigued that a cluster of pathways annotated as “T cell immunity’ were enriched in JDM muscle-only regions (Figure 2C). Scrutiny of pathways that generated this annotation revealed 22 pathways ascribed to T cells and 85 others labelled as antigen processing, regulation of immunity, MHC, or other immune cells (Table S3). Across these pathways the top DE genes were HLA (MHC) genes or those involved in MHC processing such as *TAP2* (Table S4). It is well-established that both muscle and endothelium in adult and juvenile IIM express high levels of HLA class I and II proteins which can be independent of inflammatory infiltrates^17–20^. Therefore, we ascribe this result to expression of HLA genes (driven by IFN) in muscle fibres or endothelium.

To determine whether differentially expressed pathways were upregulated or downregulated in JDM against controls, gene set enrichment analysis was performed using the fGSEA method^21^. This analysis provides complementary insights by considering the entire ranked gene list, identifying pathways with subtle but coordinated expression changes. Using fGSEA, 79 and 269 significant pathways (adjusted p value ≤ 0.05) were identified with the GO CC and GO BP gene sets, respectively, many of which overlapped with the ORA results. Full lists of the significant pathways identified via fGSEA and their associated fold changes and clusters can be found in Table S5 (GO CC) and Table S6 (GO BP).

Clustering of the 79 pathways (GO CC) using aPEAR validated the previous results, with the generated network plots reinforcing involvement of ER to Golgi transport pathways (upregulated) and mitochondrial-related pathways (downregulated, Figure S3A and Table S5). In contrast, clustering of the 269 GO BP pathways highlighted the significant upregulation of interferon signalling pathways, which fall under the cluster of “regulation of response to external stimulus” (Figure S3B and Table S6). Figure S3C shows the 15 most downregulated and 15 most upregulated pathways from both GO CC and GO BP analyses, visualised by comparing normalised enrichment scores.

### Interferon-driven signature is elevated across tissue cell types analysed in JDM patients

We next used our previously validated 15-gene IFN-stimulated gene signature^3^ to visualize variations in the expression patterns of IFN-stimulated genes across the different ROIs, from patients and controls^22^. All 15 of these genes are associated with both Type I and Type II interferon signalling as annotated in the Interferome database^23^. Heatmap visualization of normalized, scaled expression levels across ROIs revealed distinct patterns of interferon-stimulated gene (ISG) expression between patients, with unsupervised clustering of ROIs primarily by patient. One JDM patient (JDM3) exhibited lower ISG expression compared to the other two JDM patients (Figure 3A). Notably, this patient was less weak at time of biopsy than patients JDM1 and JDM2 (Table 1).

**Figure 3.**
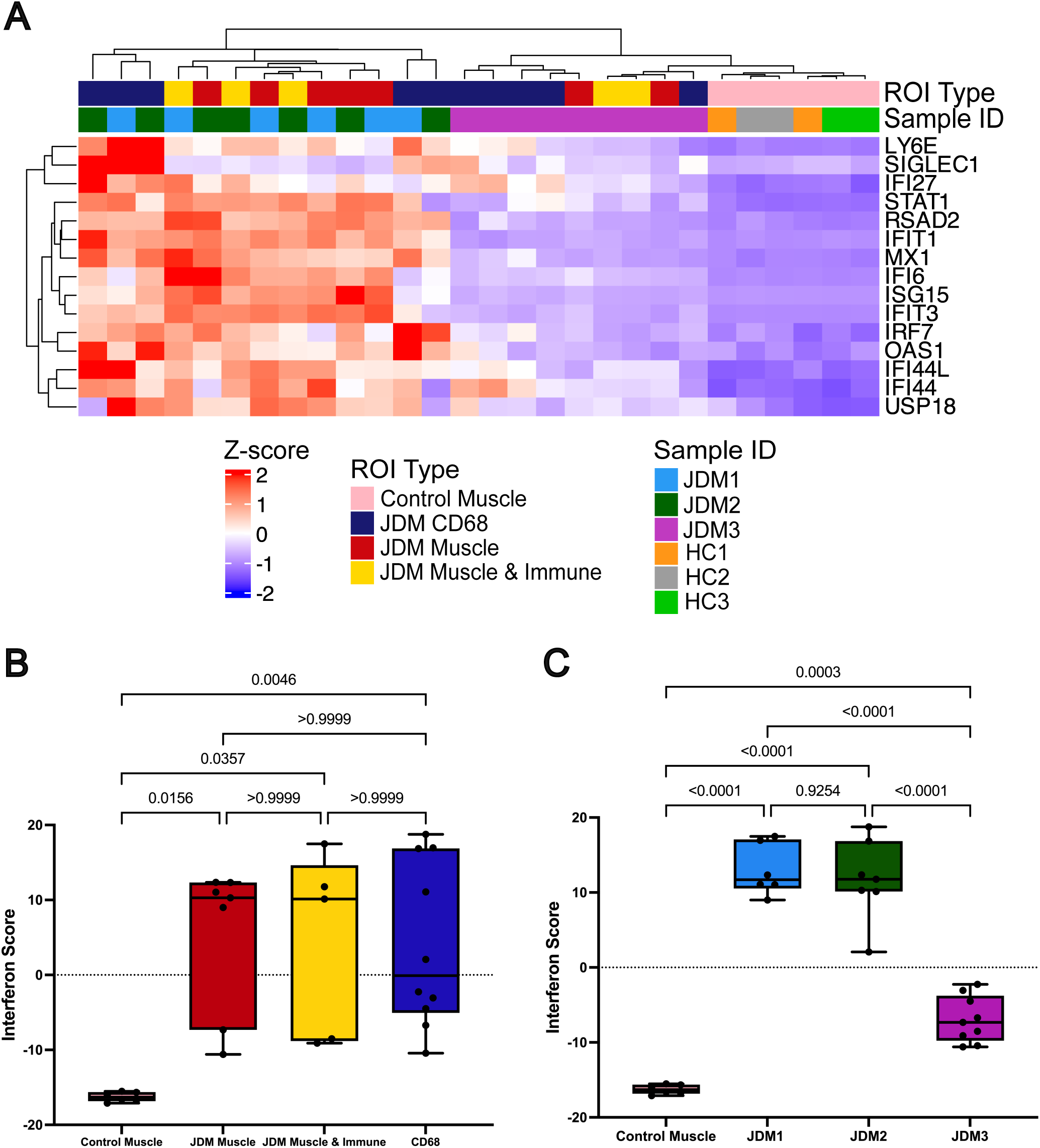
Interferon-driven signature is elevated across each cell type analysed in JDM tissue. (A) Heatmap showing the normalized, scaled expression levels of the 15-gene interferon score across the ROIs. Distinct ROI types (top row) are allocated colours pink, blue, red and yellow as shown; individual patients (second row sample ID) are allocated colours as shown. Unsupervised clustering groups the ROIs into three distinct clusters: one including ROIs patient JDM1 (blue ID) and JDM2 (green ID), a second cluster for patient JDM3 (pink ID), and a third for control ROIs (orange, grey and pale green IDs). The coloured scaled bar red to blue highlights the Z-score ranges, with red indicating a positive Z-score and blue indicating a negative Z-score. (B-C) Boxplots showing the 15-gene interferon score, calculated as the sum of Z-scores for the 15 ISGs, compared across: (B) different ROI types and (C) individual patients and controls. For (B) and (C) statistical differences between the groups were assessed using the Kruskal-Wallis test, followed by post hoc pairwise comparisons using Dunn’s test.

Quantification of the 15-gene ISG score (IFN score) demonstrated significant elevation in all ROI types from JDM tissues compared to controls. However, pairwise comparisons revealed no statistically significant differences between different ROI types within the JDM group, indicating a consistent IFN activation signature across muscle cells, immune-infiltrated regions, and CD68+ macrophage-enriched regions (Figure 3B). Analysis of IFN score values by ROI confirmed that all 3 patients had significantly higher IFN scores than controls, but that the IFN score of JDM3 was significantly lower than scores of JDM1 or JDM2 (Figure 3C).

### Mitochondrial dysfunction is consistent across all JDM patients

To further analyse mitochondrial involvement within JDM muscle tissue, the 448 DEG identified in JDM muscle ROIs compared to controls (Figure 2A) were further examined based on their associated GO terms. Of these 448 genes, 75 were annotated with the GO term “mitochondrion”. This list was carefully curated, removing known ISGs, including 5 which were part of our IFN score, and other genes which are not specific to mitochondrial function, or only interact with mitochondria under specific conditions (see Supplementary Methods). Unsupervised clustering of the 41 remaining genes revealed clusters by ROI type (and therefore cell type), rather than by patient, suggesting that mitochondrial gene expression patterns are shaped by cell type and disease state, but were largely consistent across patients (Figure 4A). Genes included in analysis shown in Figure 4 are listed in Table S7. Quantification of the mitochondrial gene score using these 41 genes demonstrated significant differences between all JDM ROIs and controls (Figure 4B). Among the JDM patient data, CD68+ macrophage regions exhibited significantly distinct mitochondrial scores compared to both JDM muscle (p < 0.0001) and control muscle ROIs (p < 0.0001) indicating cell-specific variation in mitochondrial gene expression patterns (Figure 4B), unlike the uniformity across cell types seen in interferon dysregulation in JDM (Figure 3B). No statistically significant difference was observed between the muscle and muscle+immune-cell JDM mitochondrial scores (Figure 4B, p = 0.1438). Analysis of the 41-gene score across patients confirmed that each patient had a highly significant altered mitochondrial score compared to controls, but there were no significant differences in mitochondrial scores between patients (Figure 4C). Together these results suggested that while the degree of IFN-driven pathology varied between cases, the mitochondrial abnormality was observed consistently across these 3 cases.

**Figure 4.**
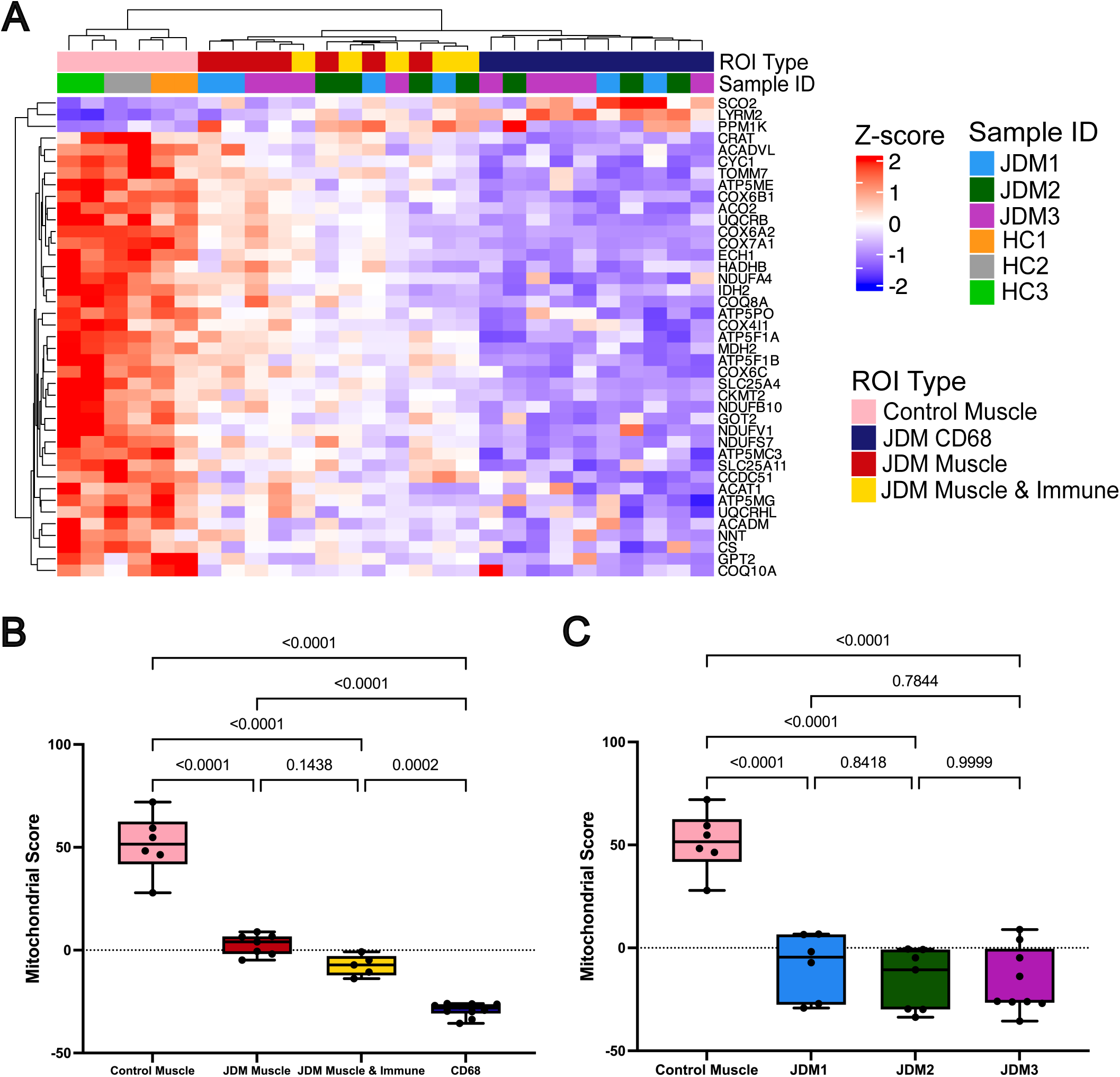
Abnormal mitochondrial signature is consistent across all JDM patients but distinct between cell types. (A) Heatmap showing the normalized, scaled expression levels of the 41-gene mitochondrial score differentially expressed in JDM muscle against control ROIs. Distinct ROI types (top row) are allocated colours pink, blue, red and yellow as shown; individual patients (second row sample ID) are allocated colours as shown. All ROI types are included in the heatmap analysis. Unsupervised clustering groups the ROIs into three distinct clusters: one including JDM muscle ROIs and muscle + immune cell ROIs, a second for CD68+ ROIs, and a third for control muscle ROIs. The coloured scaled bar highlights the Z-score ranges, with red indicating a positive Z-score and blue indicating a negative Z-score. (B-C) Boxplots of 41-gene mitochondrial scores, calculated as the sum of Z-scores for the 41 mitochondrial genes, compared across: (B) different ROI types and (C) individual patients and controls. For (B), statistical differences were assessed using one-way ANOVA, followed by post hoc pairwise comparisons using the Tukey’s HSD test. For (C), statistical differences were assessed using the Kruskal-Wallis test, followed by post hoc pairwise comparisons using Dunn’s test.

To validate these findings in an independent cohort, fGSEA analysis was performed using a publicly available RNAseq dataset, comparing four JDM with five control muscle biopsies^24^. As these data were bulk RNAseq, analysis was performed at the whole-biopsy level but not for specific regions or cell types within each biopsy. Analysis confirmed upregulation of IFN-driven and immune activation pathways, and downregulation of mitochondrial pathways (including oxidative phosphorylation, respiratory electron transport chain, ATP synthesis coupled electron transport), Figure S4A. This dataset also enabled transcriptome-wide quantification, including mitochondrial-encoded gene transcripts not captured by the GeoMx whole transcriptome atlas. Twenty of the 37 mitochondrial-encoded genes were significantly (adjusted p-value < 0.05, |Log2FC| ≥ 0.58) downregulated in JDM muscle compared to controls (Figure S4B), and this was consistent across all cases (Figure S4C). Furthermore, 10 of the 41 mitochondrial score genes were significantly and consistently downregulated in JDM in the replication cohort (Figure 4SD, 4SE), while 13 genes of the 15-gene IFN score were significantly upregulated in this cohort (Figure 4SF). Again, we observed clear patient heterogeneity in levels of IFN-driven gene expression (Figure 4SG).

### Relationship between IFN-driven and mitochondrial abnormalities in JDM Muscle

Given our intriguing finding of lack of correlation between IFN and mitochondrial pathological gene expression, we investigated this at protein level using conventional immunohistochemistry and chemical stains in a larger cohort (n=19), including 16 further UK-JDCBS JDM cases and the index 3 JDM cases, all treatment-naïve. Sections were stained and scored for IFN-driven MxA protein expression (Figure 5A-D) and mitochondrial abnormality (Figure 5E-J) (see Supplementary Methods). JDM cases exhibited a range of mitochondrial deficiency severity (6 severe, 10 mild, 3 none) and IFN-driven MxA protein expression. However, there was no correlation between these two features (Figure 5K, p=0.657). In contrast, level of MxA protein upregulation on muscle fibres correlated with clinical weakness measured by MMT8 (p=0.0369) and CMAS (p=0.0084), replicating our previous findings in 103 cases^25^.

**Figure 5.**
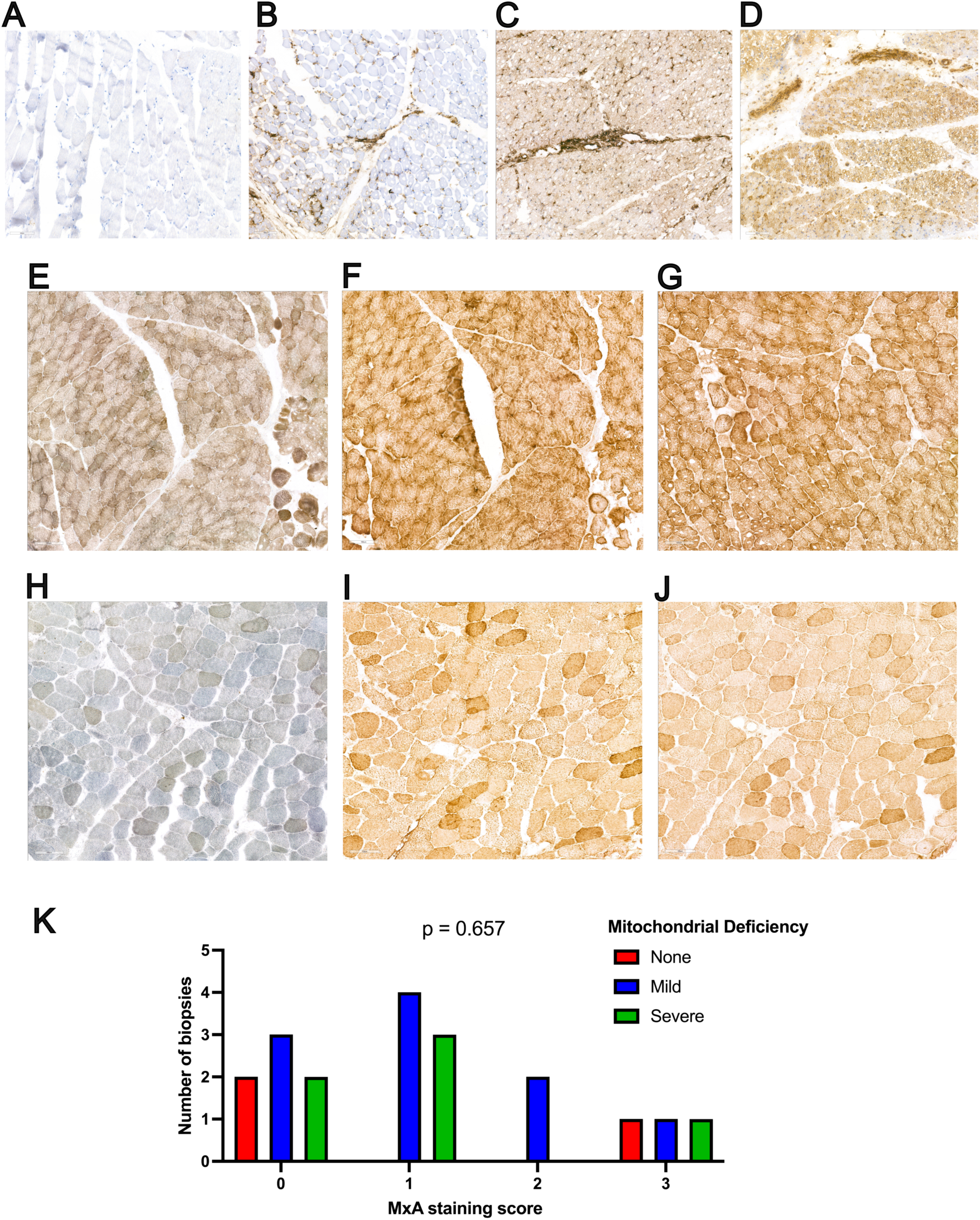
Lack of association between interferon dysregulation and mitochondrial deficiency assessed at protein level in an independent JDM cohort. (A–D) Representative immunohistochemical staining for myxovirus-resistance protein A (MxA), used to assess interferon-driven protein expression in muscle fibres. Scoring was performed using a 4-point scale: 0 = no staining (A), 1 = weak (B), 2 = moderate (C), and 3 = strong (D). (E-J) Representative staining for mitochondrial deficiency, assessed using combined COX–SDH histochemistry and supported by immunohistochemistry for MTCO1 (complex IV) and NDUFB8 (complex I). Cases were categorized as having no (0), mild (1), or severe (2) mitochondrial deficiency. (E–G) Representative staining from a JDM biopsy with mild mitochondrial deficiency: (E) COX–SDH staining, (F) MTCO1, and (G) NDUFB8. (H–J) Representative staining from a JDM biopsy with severe mitochondrial deficiency: (H) COX–SDH, (I) MTCO1, and (J) NDUFB8. (K) Summary bar plot of 19 JDM muscle biopsies, showing the distribution of mitochondrial deficiency scores (0 = none, 1 = mild, 2 = severe) across each level of interferon activity (MxA score 0–3). Bar colours indicate the level of mitochondrial deficiency (none, mild, severe) as shown. Statistical analysis was performed using the *U*-statistic permutation test.

The histological feature of perifascicular atrophy (PFA) is well-recognised in dermatomyositis and has been shown to correlate with COX deficiency and mitochondrial abnormalities in JDM and adult DM^26^. We tested whether PFA within the ROIs analysed by GeoMx was associated with mitochondrial abnormality. All 3 JDM cases demonstrated evidence of PFA. Of the 22 ROIs, 18 contained the edge of a fascicle allowing analysis for PFA (Figure S5A). PFA was present in 55% (10/18) of these ROI including regions from all 3 cases. In this small sample, no correlation was observed between PFA and mitochondrial abnormality, assessed in the 8 muscle/muscle+immune ROIs (Figure S5B). However, this result may relate to small sample size, the fact that GeoMx does not measure expression of mitochondrial-encoded genes, or that each ROI covers areas with PFA and others without, while GeoMx generates a bulk-like average 41-gene mitochondrial score for the whole ROI.

To define other biological pathways associated with mitochondrial dysfunction, two complementary approaches were performed. First, for each pathway cluster shown in Figure S3A-B, the most commonly-shared genes within each cluster were used to generate cluster-specific expression scores. In the second approach, fGSEA was run using MSigDB Hallmark gene sets, and scores calculated from the leading-edge genes of each pathway (see Supplementary Methods). Across both methods, lower mitochondrial gene expression was consistently associated with reduced activity in metabolic pathways, including oxidative phosphorylation, fatty acid metabolism, glycolysis and genes involved in reactive oxygen species (ROS) response as expected (Figure 6A-B). In addition, decreased expression of peroxisome-related genes, and downregulation of sarcomere organisation, were observed to be correlated with more severe mitochondrial abnormalities.

**Figure 6.**
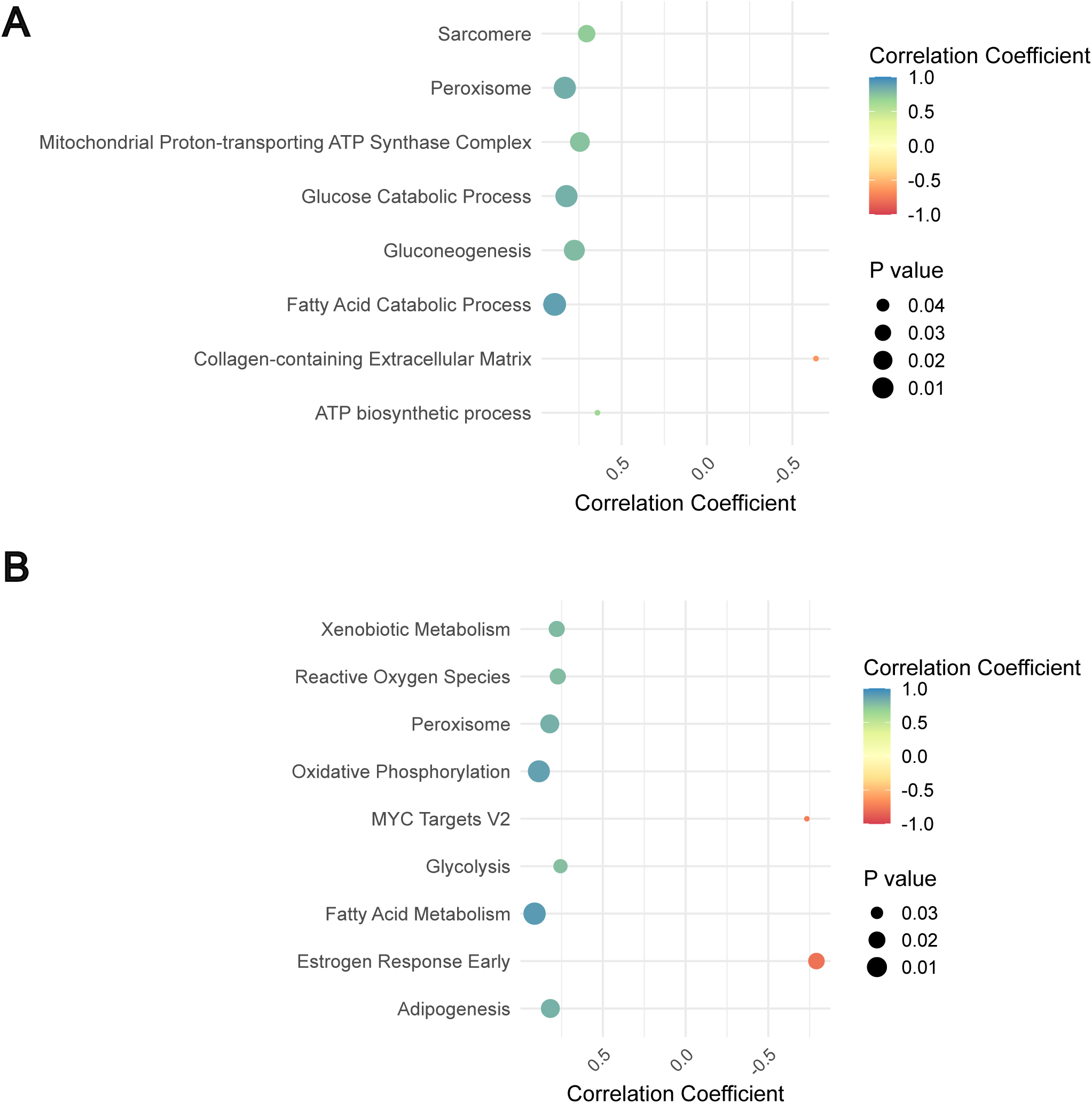
Pathway-level transcriptional signatures associated with mitochondrial dysfunction in JDM muscle. Dot plots show the correlation between the mitochondrial gene score and pathway-specific expression scores across muscle regions of interest (ROIs) from JDM biopsies. The mitochondrial gene score was calculated per ROI based on z-scores of 41 curated mitochondrial genes, using the mean and standard deviation from control muscle as a reference. (A) Correlation with GO-based pathway clusters, derived from fGSEA of JDM ROIs compared to controls. For each cluster, a core gene signature was defined by selecting genes that appeared in the leading edge of ≥ 75% of constituent pathways (B) Correlation with significantly enriched MSigDB Hallmark pathways (adjusted p ≤ 0.05) identified through gene set enrichment analysis of JDM versus control muscle. For both (A) and (B), each dot represents a single pathway or cluster. Dot size reflects the strength of the correlation (r coefficient), and dot colour indicates the statistical significance of the correlation, with deeper red shades corresponding to smaller adjusted p-values. The colour scale bar reflects the adjusted p-value.

### Transcriptomic differences between distinct regions of affected muscle and between patients

Since we observed differences in IFN-driven signature between cases, we tested for other transcriptomic differences across patients, comparing patient JDM3 with JDM1 and JDM2. Initially, muscle and muscle-immune ROIs were combined to increase statistical power. Analysis comparing JDM3 vs. JDM1+JDM2 revealed 176 DEGs between groups (100 upregulated, 76 downregulated genes in patients JDM1+JDM2, Figure S6A). Pathway analysis indicated 208 enriched GO BP pathways; cluster network analysis highlighted interferon and immune-related pathway clusters, including “positive regulation of type I interferon production” (confirming our 15-gene IFN-score result, Figure 3), “antigen processing and presentation via MHC Class I via ER pathways”, and “antibacterial humoral response” (Figure S6B). In this comparison between patients, mitochondrial pathways were not differentially expressed, confirming earlier results suggesting consistent mitochondrial abnormalities in all cases. Separate DGE analysis performed between the CD68+ cell ROIs of patient JDM3 against those of patients JDM1+JDM2 revealed 63 DEGs, (43 upregulated, 20 downregulated in JDM1+JDM2, Figure S7A). Within this analysis, 148 GO BP pathways were significantly enriched, with the most prominent cluster being “negative regulation of viral genome replication” (Figure 7B), indicating that expression of ISGs in muscle-infiltrating CD68+ cells also differed between the JDM patients. We also conducted DGE analysis for each ROI type comparing JDM1+JDM3 vs JDM2, and JDM2+JDM3 vs JDM1. Those comparisons did not reveal meaningful differences, yielding between 0 to 4 DEGs only (data not shown)

**Figure 7.**
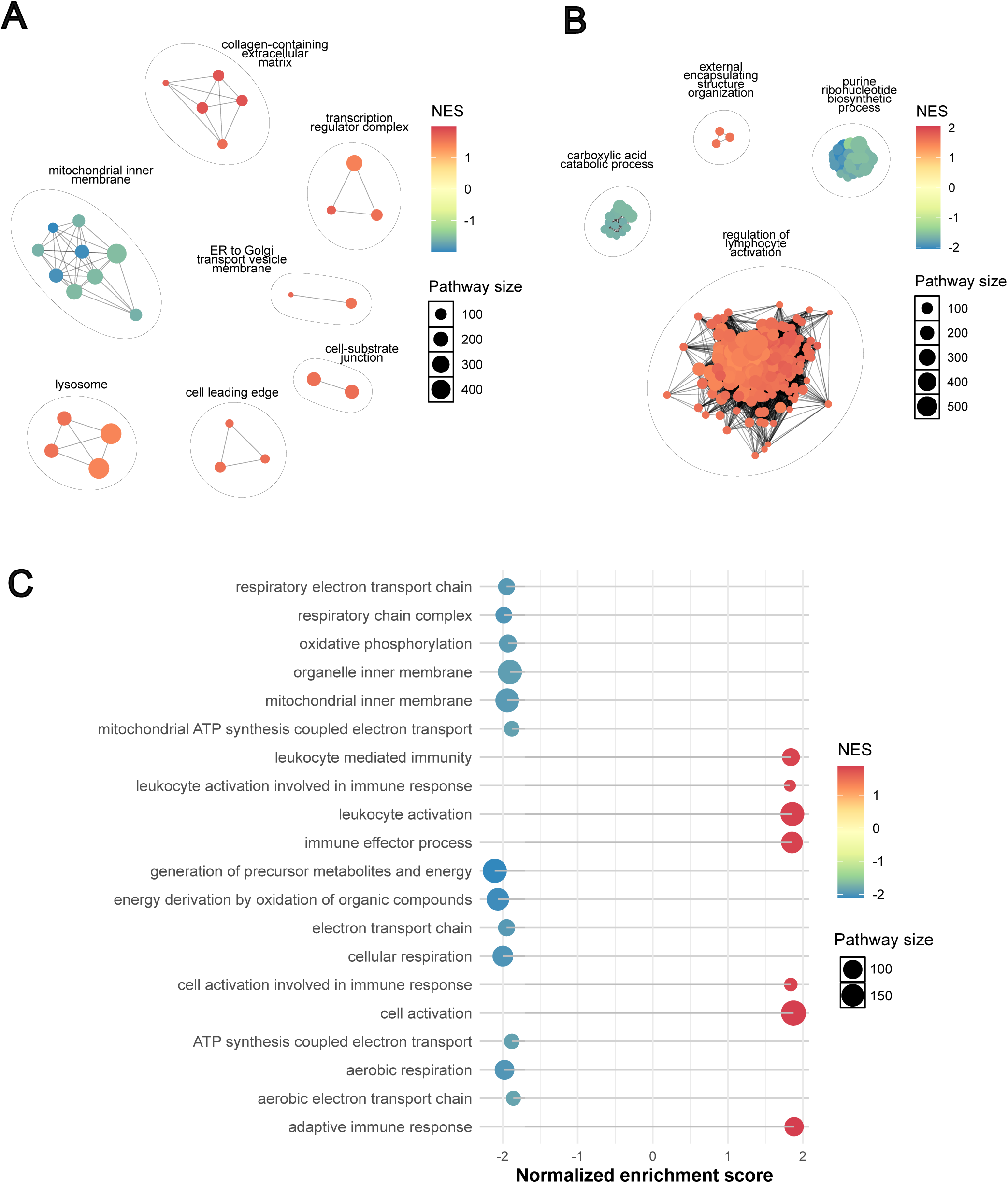
Gene set enrichment analysis reveals increased immune pathways and decreased mitochondrial functional pathways in JDM immune cell-rich muscle regions compared to JDM muscle only regions. (A-B) Cluster plots of enriched Gene Ontology pathways, based on fGSEA performed on all genes for each comparison ranked by p-value and fold change. (A) Plot of enriched GO Cellular Component pathways. (B) Plot of enriched GO Biological Process pathways. (C) Dot plot of the 20 GO BP and CC pathways with the highest absolute normalized enrichment score (NES). In panels (A-C), larger dots represent pathways associated with a higher number of genes (shown in key, pathway size), and the dot colour reflects the NES, with blue indicating negative NES and red indicating positive (shown in colour bar, NES).

Finally, we explored transcriptional patterns in muscle-immune ROI (which had dense infiltrates of CD45+ immune cells, including CD3+/CD68+/CD20+ cells), compared to muscle-fibre only regions. Initial DGE analysis using the limma-voom pipeline suggested no significant DEGs between these ROI groups, based on the same fold change and adjusted p-value criteria (data not shown). However, fGSEA analysis revealed 28 significant pathways in the GO CC gene set and 317 significant pathways in the GO BP gene set (Figure 7A, 7B respectively). These pathway-level findings suggested that immune-infiltrated muscle regions exhibited further downregulation of mitochondrial and metabolic pathways, alongside increased expression of immune-related and leukocyte activation pathways (Figure 7C). While not captured at the significant DEG level, these transcriptional differences are consistent with the expected biological impact of immune infiltration. These significant pathways are found in Tables S8 (GO CC) and S9 (GO BP).

## DISCUSSION

Our previous work and that of others studying affected muscle in JDM, a key target tissue of pathology, has demonstrated upregulated expression of IFN-driven gene and protein expression. In muscle fibres this includes overexpression of class I MHC protein (also known as HLA class I)^6, 20, 27^, and myxovirus-resistance protein A (MxA1)^25^. Expression of IFN-driven proteins is variable between cases, but typically correlates with degree of weakness^25^. Many studies have confirmed a strong IFN-driven transcriptional signature in both muscle and blood of both adult and paediatric patients with IIM^3, 28–31^. Therapies which target IFN or IFN-signalling are increasingly being tested in myositis^32^.

Our previous gene expression analysis of peripheral blood lineage-sorted immune cells from patients with JDM revealed not only this known IFN-driven signature, but also a signature of mitochondrial dysfunction, most marked in blood monocytes^3^ which was confirmed at the functional level and found to persist, and was not restored to normal by treatment, despite clear reduction in expression of ISGs on treatment. Furthermore, we showed that mitochondrial DNA was abnormally released in patient monocytes and could drive production of IFN and downstream ISGs, an effect that was mediated via C-GAS-STING and TLR9 pathways.

To date the majority of transcriptional studies published using muscle tissue of patients with IIM have used bulk RNA sequencing. In these datasets it is challenging to understand or define the role of different cells, or to identify how specific tissue micro-niches contribute to pathology. In this study, we have for the first time used spatial transcriptomic methods to analyse muscle tissue from children with JDM obtained at diagnosis prior to starting treatment compared to age-matched healthy muscle. Thus, our data are not confounded by effects of medication. We used Nanostring GeoMx® DSP to generate transcriptional data of three specific region types within the muscle. This allowed comparison of regions within the disease tissue including those where infiltrating immune cells are scarce, or not detected, in regions of dense clusters of inflammatory cells, and in regions where CD68+ infiltrating macrophages were predominant. Our data confirm successful use of this platform for analysis using historical cryopreserved muscle tissue.

Initial analysis of JDM compared to control tissue confirmed high expression of IFN-stimulated genes in all regions analysed. Furthermore, strong upregulation of pathways related to protein transport through the ER and Golgi, and ER stress, was observed, aligning well with our previous studies in human muscle and a transgenic mouse myositis model^4, 33, 34^. We also demonstrate that, as previously observed in blood monocytes, in the JDM tissue itself, muscle fibres have a highly dysregulated mitochondrial signature. Dysregulated pathways included those related to mitochondrial function, respiratory chain, and mitochondrial proton transport: their expression was downregulated in muscle. In addition, the signal for mitochondrial and respiratory chain related genes was distinct in CD68-enriched regions. By analysing RNAseq data from a second JDM cohort, we validated our results for these nuclear-encoded mitochondrial genes and additionally showed downregulation of mitochondrial-encoded genes. We investigated the relationship of the signatures to clinical activity as assessed by muscle weakness. We observed a correlation between IFN signature and weakness, whereby one patient who was minimally weak (MMT8 score 78) had lower expression of IFN-induced signature, than the two patients who were significantly weak (MMT8 scores 44 and 42). Interestingly this was different for mitochondrial signature detected in tissue, which was independent of the IFN signature. The downregulation of mitochondrial pathways was consistently observed for the index three patients, regardless of muscle strength, and across the three patients this result showed minimal variance for the 41-gene mitochondrial score we defined. We extended these findings in 19 cases at protein level, and replicated the observation of no correlation between level of IFN-driven protein expression with mitochondrial abnormality. A recent study suggested that IFNγ itself leads to mitochondrial dysfunction and oxidative stress in both a mouse model and patients with adult IIM^10^. In contrast, our previous work in JDM monocytes suggested that circulating oxidised mitochondrial DNA itself may activate IFN pathways. The current study suggests that mitochondrial dysfunction can be equally prominent in patients with either low or high IFN score at both RNA and protein level, implying that IFN-independent drivers of mitochondrial dysfunction exist. If so, blocking of IFN by new therapeutic agents may not adequately control all aspects of disease pathology. Supporting this, recent evidence in IBM and PM suggests that mitochondrial-encoded gene mutations lead to inflammation and immune-cell infiltration, activating the cGAS/STING pathway, in the absence of IFN^35^. In a model of Parkinsons disease (PD) and PD patients, mitochondrial DNA damage triggers neuron pathology, which is, remarkably, exacerbated in the IFN-deficient mouse^36^. Collectively these findings suggest that intersection between mitochondrial biology and interferon is bidirectional and complex, and that pathologies driven by altered mitochondrial DNA and function can act independently of IFN. Our demonstration that mitochondrial dysfunction correlates with altered function of the peroxisome, an organelle key to metabolic pathways which communicates with mitochondria may reveal further potential novel treatment targets^37^.

Our analysis of JDM muscle compared to healthy muscle also revealed other dysregulated pathways associated with disease, including ER to Golgi transport, sarcomere and T cell activation. We attribute this to the highly upregulated HLA class I and II DEG which are known to be upregulated in muscle fibres themselves in JDM.

Direct comparison of muscle-only to muscle+immune regions revealed pronounced downregulation of mitochondrial and metabolic pathways, alongside increased expression of immune-related pathways, in immune-infiltrated regions. This indicates that relative mitochondrial abnormalities differ between cell types in inflamed tissue, and concurs with evidence that the mitochondrial transcriptome differs across tissue types^38^. To precisely dissect specific tissue niches and cell interactions in JDM, future single-cell spatial analyses will provide more precise understanding of cell-specific contributions and define cell-cell interactions in JDM muscle tissue. Single cell spatial analysis of JDM muscle is underway to allow us to distinguish these issues.

This study has several limitations. The initial sample size (n=3 in each group) was small which limited our ability to detect gene expression differences with a small effect size, which may still be important to pathology: therefore, our findings should be interpreted with caution, especially when comparing 2 patients to one other. Despite this, key significant differences between patients and controls were replicated in a second cohort and our findings were validated at protein level in a larger cohort. Our previous analysis of 103 JDM muscle biopsies showed significant histological heterogeneity across cases which varied in part by MSA status^25^. The current study was not powered to analyse for association between specific MSA status and transcriptional differences. A larger cohort will therefore be essential to capture the full variance in patient phenotype, MSA status, clinical severity and pathology, to provide a robust understanding of JDM-associated gene expression changes. Furthermore analysis of tissue-infiltrating CD68+ macrophages lacked a control comparison group due to the absence of CD68+ cells in normal muscle tissue limiting our ability to identify the unique transcriptomic signature of infiltrating macrophages against healthy macrophages.

This study represents the first application of spatial transcriptomics to JDM patient muscle biopsies using the GeoMx® technology, unveiling critical insights into mitochondrial and immune dysregulation at the tissue level. Importantly, we have defined parallel gene expression patterns in both the muscle and macrophage-rich regions of JDM muscle, as previously identified in blood monocytes. Together these data suggest that mitochondrial dysfunction is present across tissues and, that within the muscle itself this signal is readily detectable in both CD68+ infiltrating cells and muscle fibres. Our results also indicate that this mitochondrial dysfunction is uncoupled from the characteristic IFN-driven signature. Given that almost 50% of patients with JDM are not well-controlled on standard first-line treatments^2^, which are currently focused on the suppression of immune pathways, our demonstration that mitochondrial dysfunction is present in muscle of the majority of cases of JDM in this study has important implications. First, adjunctive therapies which target the mitochondrial pathology could be tested in combination with current treatments. Secondly, this signature may be readily tested and used to drive treatment choices in the future. Future work to define whether the mitochondrial signature in tissue correlates with that in blood will be important for design of stratification tools in trials of novel agents targeting the mitochondrial abnormality.

Our findings advance the understanding of JDM pathogenesis and lay the groundwork for future research into the use of therapies which specifically target mitochondrial dysfunction and immune-mediated damage.

### Patient and public involvement and engagement

Patients and carers were involved at every stage of this research, including study conception, design, delivery, and analysis, through our partnership stakeholder group, the JDCBS PPIE group. The study was discussed at our annual JDCBS study day and also at the UK JDM Family Day, where families, patients and parents provided input. All patients and families who are part of the study across the UK hear about updates via a regular newsletter as well as on our dedicated <u>website</u>.

## Acknowledgments

We thank all of the families, patients, parents and carers who contributed to the study and allowed us to use samples and data for this work. We are very grateful to Mr Dario Cancemi for excellent Study management, to the clinical team who assist with data collection, to Dr Lizzy Rosser for critical review and to the BRAIN Bank UK for access to control cases. We are grateful to the members of the <u>JDCBS research group (JDRG)</u> and who are listed in the supplementary materials. We thank the team at Nanostring for valuable guidance, the GOSH histopathology team including Abigail White and Mwaka Mshanga, and Darren Chambers, Department of Neuropathology, UCL Queen Square Institute of Neurology for chemical stains.

The data presented in this manuscript have not been published previously. However, some of the data were submitted as an abstract for the European Alliance of Associations for Rheumatology (EULAR) 2025 congress.

## Contributions

AES contributed to data curation, led the development of the formal analysis and methodology pipeline, visualization, and drafting of the original manuscript. MK, LN, NMLE and MCB contributed to data curation and methodology development. APC, CBM and SE contributed to analysis. ChP, MO, ClP and YG contributed to data collection and curation. AM and OO were responsible for histopathology analysis. LRW and MGLW conceptualized the study, acquired funding and co-led analysis, methodology development, project supervision. LRW, MGLW and MK contributed to both original drafting and editing of the manuscript. All authors reviewed and approved the final manuscript for submission.

## Supplementary

see Supplementary materials.

## Data availability statement

The data that support the findings of this study are available from the corresponding authors upon reasonable request. Script used in analyses is openly available in Github (https://github.com/WedderburnLab/GeoMx-pipeline).

## REFERENCES

1. Kim H. Updates on interferon in juvenile dermatomyositis: pathogenesis and therapy. Curr Opin Rheumatol 2021;33(5):371–77. doi: 10.1097/bor.0000000000000816

2. Papadopoulou C, Chew C, Wilkinson MGL, et al. Juvenile idiopathic inflammatory myositis: an update on pathophysiology and clinical care. Nat Rev Rheumatol 2023;19(6):343–62. doi: 10.1038/s41584-023-00967-9 [published Online First: 20230515]

3. Wilkinson MGL, Moulding D, McDonnell TCR, et al. Role of CD14+ monocyte-derived oxidised mitochondrial DNA in the inflammatory interferon type 1 signature in juvenile dermatomyositis. Ann Rheum Dis 2023;82(5):658–69. doi: 10.1136/ard-2022-223469 [published Online First: 20221223]

4. Nistala K, Varsani H, Wittkowski H, et al. Myeloid related protein induces muscle derived inflammatory mediators in juvenile dermatomyositis. Arthritis Res Ther 2013;15(5):R131. doi: 10.1186/ar4311 [published Online First: 20130923]

5. Yasin SA, Schutz PW, Deakin CT, et al. Histological heterogeneity in a large clinical cohort of juvenile idiopathic inflammatory myopathy: analysis by myositis autoantibody and pathological features. Neuropathol Appl Neurobiol 2019;45(5):495–512. doi: 10.1111/nan.12528 [published Online First: 20190311]

6. Varsani H, Charman SC, Li CK, et al. Validation of a score tool for measurement of histological severity in juvenile dermatomyositis and association with clinical severity of disease. Ann Rheum Dis 2015;74(1):204–10. doi: 10.1136/annrheumdis-2013-203396 [published Online First: 20130924]

7. Arnold L, Henry A, Poron F, et al. Inflammatory monocytes recruited after skeletal muscle injury switch into antiinflammatory macrophages to support myogenesis. J Exp Med 2007;204(5):1057–69. doi: 10.1084/jem.20070075 [published Online First: 20070507]

8. Hernandez-Torres F, Matias-Valiente L, Alzas-Gomez V, et al. Macrophages in the Context of Muscle Regeneration and Duchenne Muscular Dystrophy. International Journal of Molecular Sciences 2024;25(19):10393.

9. Li Z, Liu H, Xie Q, et al. Macrophage involvement in idiopathic inflammatory myopathy: pathogenic mechanisms and therapeutic prospects. J Inflamm (Lond) 2024;21(1):48. doi: 10.1186/s12950-024-00422-w [published Online First: 20241126]

10. Abad C, Pinal-Fernandez I, Guillou C, et al. IFNγ causes mitochondrial dysfunction and oxidative stress in myositis. Nature Communications 2024;15(1):5403. doi: 10.1038/s41467-024-49460-1

11. Piñeiro AJ, Houser AE, Ji AL. Research Techniques Made Simple: Spatial Transcriptomics. J Invest Dermatol 2022;142(4):993–1001.e1. doi: 10.1016/j.jid.2021.12.014

12. Böning S, Schneider F, Huber AK, et al. Region of interest localization, tissue storage time, and antibody binding density-a technical note on the GeoMx® Digital Spatial Profiler. Immunooncol Technol 2024;23:100727. doi: 10.1016/j.iotech.2024.100727 [published Online First: 20240820]

13. Lam CG, Manlhiot C, Pullenayegum EM, et al. Efficacy of intravenous Ig therapy in juvenile dermatomyositis. Ann Rheum Dis 2011;70(12):2089–94. doi: 10.1136/ard.2011.153718 [published Online First: 20111006]

14. Varsani H, Newton KR, Li CK, et al. Quantification of normal range of inflammatory changes in morphologically normal pediatric muscle. Muscle Nerve 2008;37(2):259–61. doi: 10.1002/mus.20898

15. Law CW, Chen Y, Shi W, et al. voom: Precision weights unlock linear model analysis tools for RNA-seq read counts. Genome Biol 2014;15(2):R29. doi: 10.1186/gb-2014-15-2-r29 [published Online First: 20140203]

16. Kerseviciute I, Gordevicius J. aPEAR: an R package for autonomous visualization of pathway enrichment networks. Bioinformatics 2023;39(11) doi: 10.1093/bioinformatics/btad672

17. Emslie-Smith AM, Arahata K, Engel AG. Major histocompatibility complex class I antigen expression, immunolocalization of interferon subtypes, and T cell-mediated cytotoxicity in myopathies. Hum Pathol 1989;20(3):224–31. doi: 10.1016/0046-8177(89)90128-7

18. Englund P, Lindroos E, Nennesmo I, et al. Skeletal muscle fibers express major histocompatibility complex class II antigens independently of inflammatory infiltrates in inflammatory myopathies. Am J Pathol 2001;159(4):1263–73. doi: 10.1016/s0002-9440(10)62513-8

19. Englund P, Nennesmo I, Klareskog L, et al. Interleukin-1alpha expression in capillaries and major histocompatibility complex class I expression in type II muscle fibers from polymyositis and dermatomyositis patients: important pathogenic features independent of inflammatory cell clusters in muscle tissue. Arthritis Rheum 2002;46(4):1044–55. doi: 10.1002/art.10140

20. Li CK, Varsani H, Holton JL, et al. MHC Class I overexpression on muscles in early juvenile dermatomyositis. J Rheumatol 2004;31(3):605–9.

21. Korotkevich G, Sukhov V, Budin N, et al. Fast gene set enrichment analysis. bioRxiv 2021:060012. doi: 10.1101/060012

22. Gu Z, Eils R, Schlesner M. Complex heatmaps reveal patterns and correlations in multidimensional genomic data. Bioinformatics 2016;32(18):2847–49. doi: 10.1093/bioinformatics/btw313

23. Rusinova I, Forster S, Yu S, et al. INTERFEROME v2.0: an updated database of annotated interferon-regulated genes. Nucleic Acids Research 2012;41(D1):D1040–D46. doi: 10.1093/nar/gks1215

24. Roberson EDO, Mesa RA, Morgan GA, et al. Transcriptomes of peripheral blood mononuclear cells from juvenile dermatomyositis patients show elevated inflammation even when clinically inactive. Sci Rep 2022;12(1):275. doi: 10.1038/s41598-021-04302-8 [published Online First: 20220107]

25. Soponkanaporn S, Deakin CT, Schutz PW, et al. Expression of myxovirus-resistance protein A: a possible marker of muscle disease activity and autoantibody specificities in juvenile dermatomyositis. Neuropathol Appl Neurobiol 2019;45(4):410–20. doi: 10.1111/nan.12498 [published Online First: 20180604]

26. Hedberg-Oldfors C, Lindgren U, Visuttijai K, et al. Respiratory chain dysfunction in perifascicular muscle fibres in patients with dermatomyositis is associated with mitochondrial DNA depletion. Neuropathol Appl Neurobiol 2022;48(7):e12841. doi: 10.1111/nan.12841 [published Online First: 20220806]

27. Wedderburn LR, Varsani H, Li CK, et al. International consensus on a proposed score system for muscle biopsy evaluation in patients with juvenile dermatomyositis: a tool for potential use in clinical trials. Arthritis Rheum 2007;57(7):1192–201. doi: 10.1002/art.23012

28. Pinal-Fernandez I, Casal-Dominguez M, Derfoul A, et al. Identification of distinctive interferon gene signatures in different types of myositis. Neurology 2019;93(12):e1193–e204. doi: 10.1212/wnl.0000000000008128 [published Online First: 20190821]

29. Corman VM, Preusse C, Melchert J, et al. Deep RNA sequencing of muscle tissue reveals absence of viral signatures in dermatomyositis. Free Neuropathol 2024;5 doi: 10.17879/freeneuropathology-2024-5149 [published Online First: 20240104]

30. Bilgic H, Ytterberg SR, Amin S, et al. Interleukin-6 and type I interferon-regulated genes and chemokines mark disease activity in dermatomyositis. Arthritis Rheum 2009;60(11):3436–46. doi: 10.1002/art.24936

31. Baechler EC, Bauer JW, Slattery CA, et al. An interferon signature in the peripheral blood of dermatomyositis patients is associated with disease activity. Mol Med 2007;13(1-2):59–68. doi: 10.2119/2006-00085.Baechler

32. Ll Wilkinson MG, Deakin CT, Papadopoulou C, et al. JAK inhibitors: a potential treatment for JDM in the context of the role of interferon-driven pathology. Pediatr Rheumatol Online J 2021;19(1):146. doi: 10.1186/s12969-021-00637-8 [published Online First: 20210925]

33. Rayavarapu S, Coley W, Nagaraju K. Endoplasmic reticulum stress in skeletal muscle homeostasis and disease. Curr Rheumatol Rep 2012;14(3):238–43. doi: 10.1007/s11926-012-0247-5

34. Li CK, Knopp P, Moncrieffe H, et al. Overexpression of MHC class I heavy chain protein in young skeletal muscle leads to severe myositis: implications for juvenile myositis. Am J Pathol 2009;175(3):1030–40. doi: 10.2353/ajpath.2009.090196 [published Online First: 20090821]

35. Kleefeld F, Cross E, Lagos D, et al. Mitochondrial damage is associated with an early immune response in inclusion body myositis. Brain 2025 doi: 10.1093/brain/awaf118 [published Online First: 20250407]

36. Tresse E, Marturia-Navarro J, Sew WQG, et al. Mitochondrial DNA damage triggers spread of Parkinson’s disease-like pathology. Mol Psychiatry 2023;28(11):4902–14. doi: 10.1038/s41380-023-02251-4 [published Online First: 20231002]

37. Picca A, Faitg J, Auwerx J, et al. Mitophagy in human health, ageing and disease. Nat Metab 2023;5(12):2047–61. doi: 10.1038/s42255-023-00930-8 [published Online First: 20231130]

38. Ali AT, Boehme L, Carbajosa G, et al. Nuclear genetic regulation of the human mitochondrial transcriptome. Elife 2019;8 doi: 10.7554/eLife.41927 [published Online First: 20190218]

