## Supplementary Materials for "Spatial transcriptomic analysis of muscle biopsy from treatment-naive juvenile dermatomyositis patients reveals mitochondrial abnormalities despite disease-related interferon driven signature"

**Supplemental materials**

**List of contents**

**Supplemental Figures (S1-S7) and Figure Legends**

**Supplemental Tables and Table Legends**

**Tables S3, S5, S6, S8 and S9 are found in the file: “Supplemental_Tables_S3_S5_S6_S8_S9.xlsx”**

**Methods**

**Supplemental References**

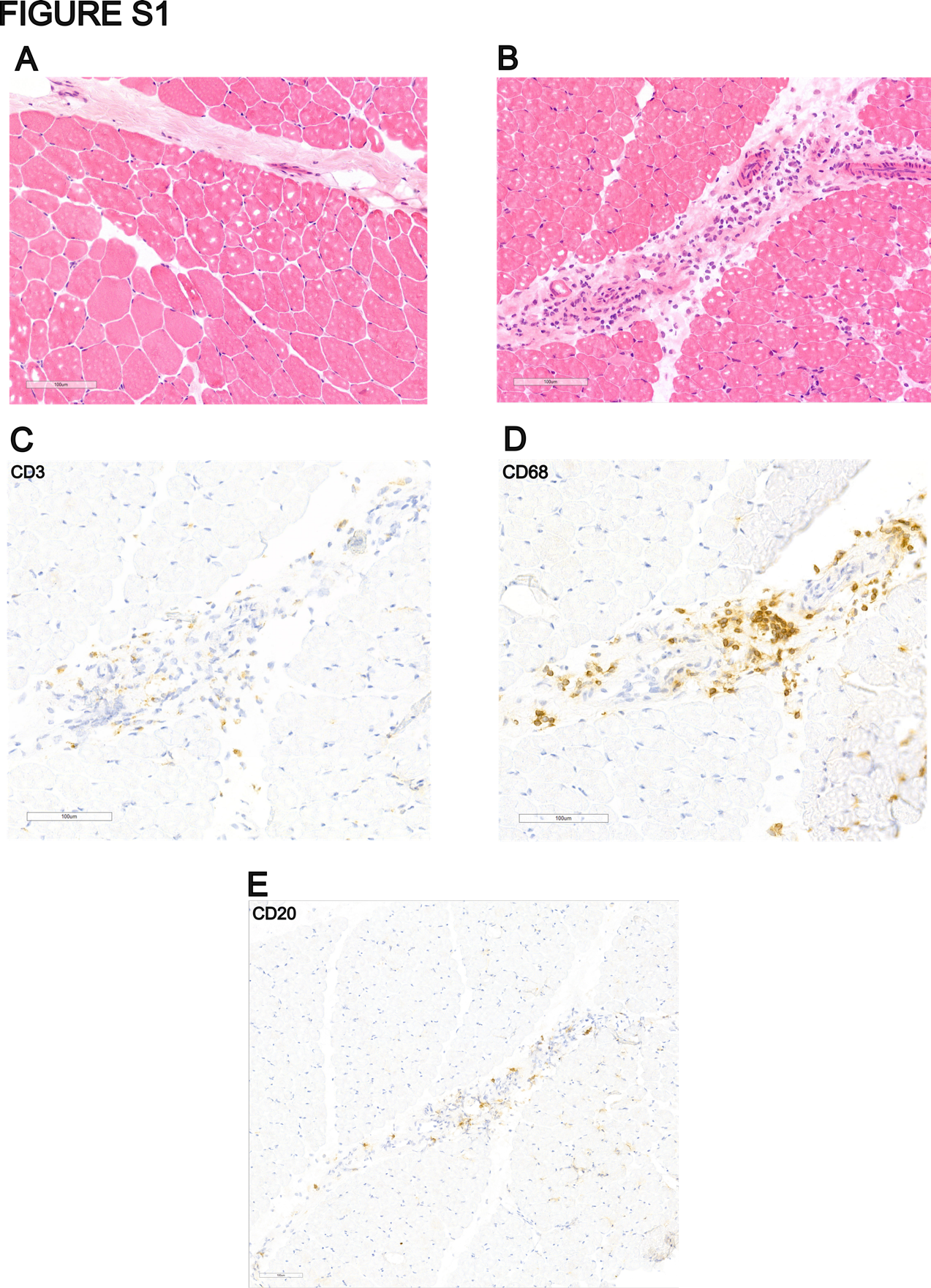

**Supplemental** **Figure S1. JDM and Control muscle biopsy representative immunohistochemistry and morphology staining.** Representative images of JDM and control muscle sections. (A) H&E staining of control muscle; (B) H&E staining of JDM muscle showing areas representative of inflammatory infiltrates; (C) CD3 staining highlighting T-lymphocyte infiltrates in JDM muscle; (D) CD68 staining showing macrophage infiltrates in JDM muscle; (E) CD20 staining highlighting B-cell infiltrates in JDM muscle.

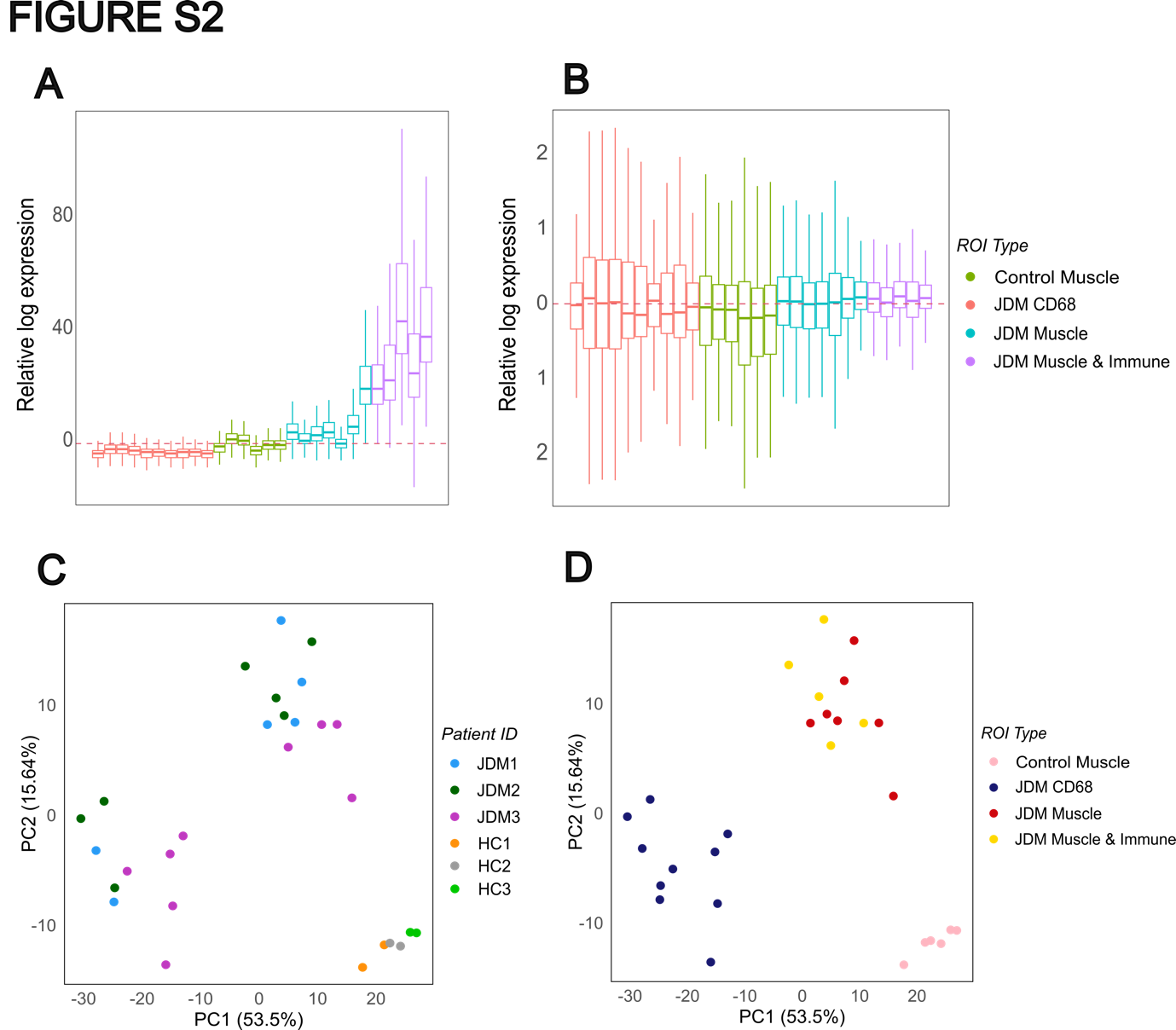

**Supplemental** **Figure S2. Relative log expression and principal component analysis of the ROIs.** (A-B) Panels displaying relative log expression values of gene expression data from the 28 patient and control ROIs, including control muscle regions (green colour), JDM regions enriched with muscle cells (cyan colour), mixed muscle and immune cells (purple colour), and CD68+ stained areas (red colour). (A) Plot showing data before normalization. (B) Plot showing values after logCPM normalization. (C-D) PCA plots representing variation across patients, controls and different ROIs. (C) PCA plot where each dot represents an ROI, coloured based on patient or control; JDM patients 1-3 coloured by cyan, green and purple symbols respectively; controls coloured by orange, grey and pale green symbols, as shown; (D) PCA plot where each dot represents an ROI and is coloured based on the type of ROI; blue CD68 regions; red JDM muscle regions; yellow JDM muscle and immune cell regions; control regions, pink symbols as shown.

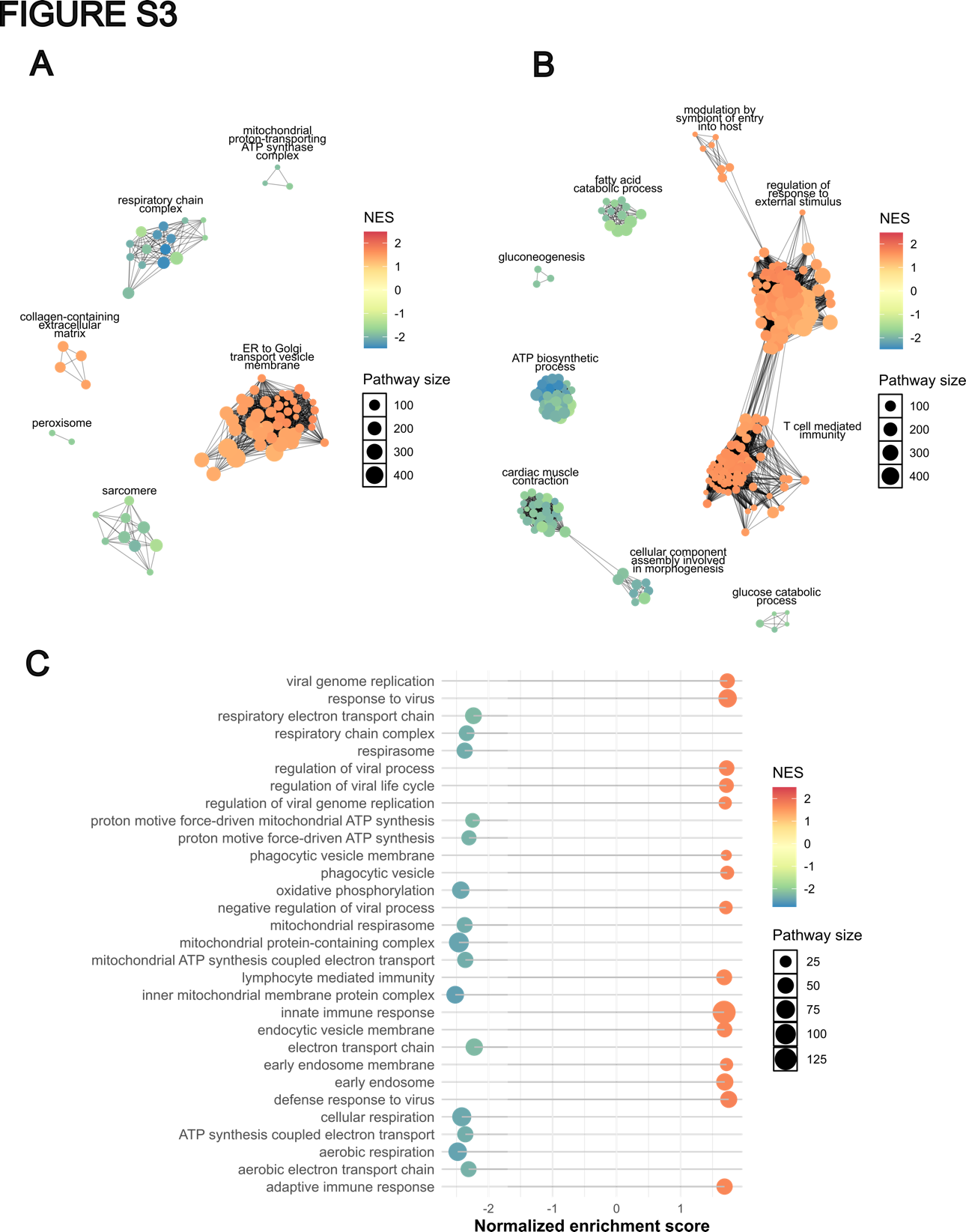

**Supplemental** **Figure S3. Gene set enrichment analysis in JDM muscle vs. Control muscle.** (A-B) aPEAR cluster plots of enriched Gene Ontology pathways, based on GSEA performed on all genes for each comparison ranked by p-value and fold change. (A) Plot of enriched GO Cellular Component pathways in JDM vs. control muscle ROIs. (B) Plot of enriched GO Biological Process pathways in JDM vs. control muscle ROIs. (C) Dot plot of the 15 GO pathways with the highest normalized enrichment score (NES) and the 15 pathways with the lowest NES. For (A-C), larger dots represent pathways associated with a higher number of genes (shown in key, pathway size), dot colour reflects NES, with blue indicating negative NES and red indicating positive (colour bar, NES).

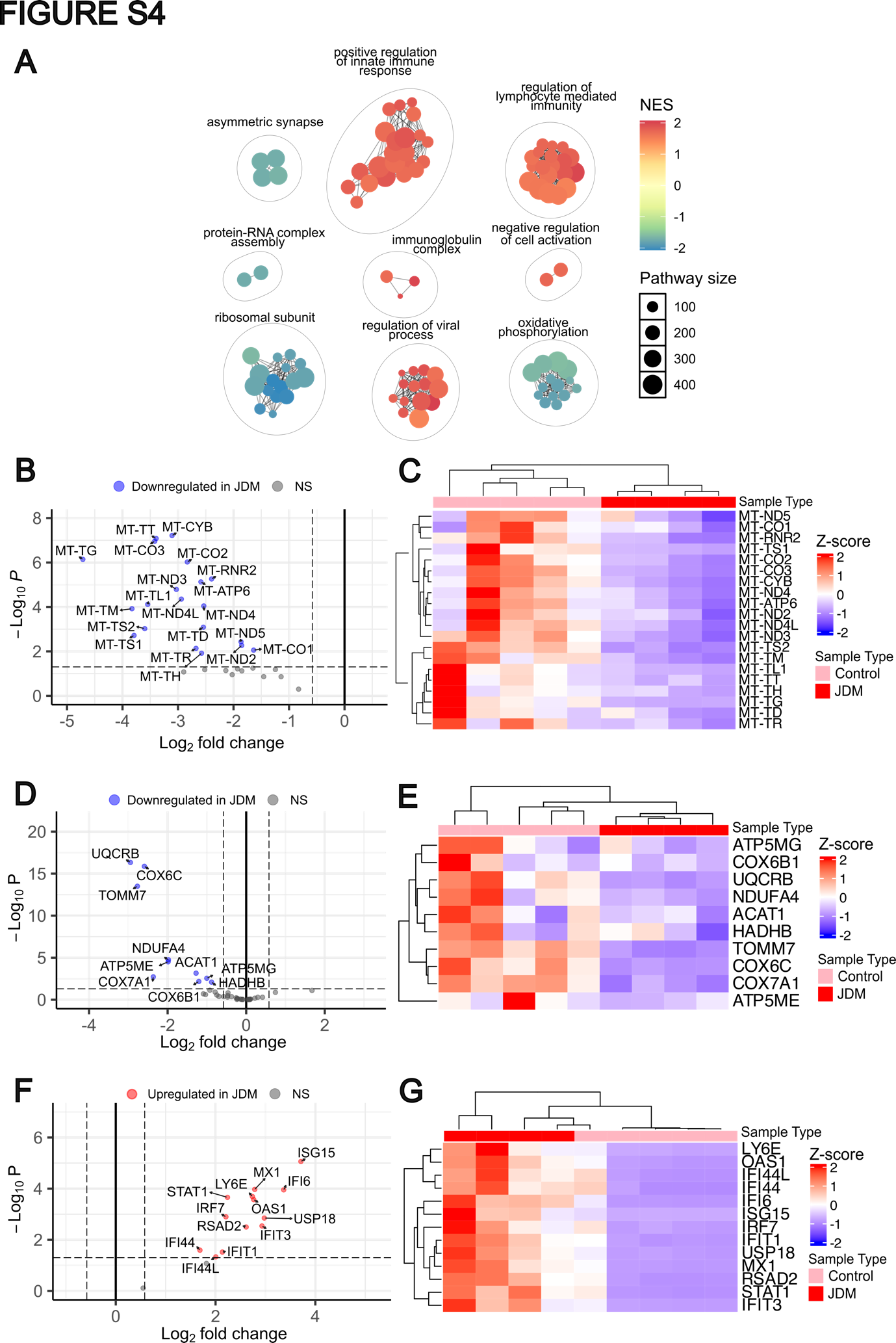

**Supplemental** **Figure S4**. **Validation of mitochondrial and IFN signatures in an independent cohort**. (A) Network plot generated using GO terms for Biological Process and Cellular Component, based on GSEA performed on all genes ranked by p-value and fold change from a validation cohort consisting of 4 JDM muscle biopsies and 5 controls. Larger dots represent pathways associated with a higher number of genes (shown in key, pathway size), dot colour reflects NES, with blue indicating negative NES and red indicating positive (colour bar, NES). (B) Volcano plot showing significant downregulation of 20 out of 37 mitochondrial-encoded genes in JDM compared to controls. (C) Heatmap of the 20 significantly differentially expressed mitochondrial-encoded genes showing distinct clustering of JDM and control samples. (D) Volcano plot showing downregulation of 10 out of 41 genes from the mitochondrial nuclear-coded score panel in JDM. (E) Heatmap of the 10 nuclear-encoded mitochondrial genes, again showing clear separation between JDM and control groups. (F) Volcano plot demonstrating upregulation of 13 out of 15 interferon-stimulated genes (ISGs) in the validation cohort. (G) Heatmap of the 13 ISGs showing distinct clustering of JDM and controls. In panel (A) larger dots represent pathways associated with a higher number of genes (shown in key, pathway size), and the dot colour reflects the NES, with blue indicating negative NES and red indicating positive (shown in colour bar, NES). For panels (B), (D) and (F), genes with an adjusted p-value ≤ 0.05 and |log2FC| ≥ 0.58 are considered statistically significant. Red points represent significantly upregulated genes, blue significantly downregulated genes. NS = not significant; FC = fold change.

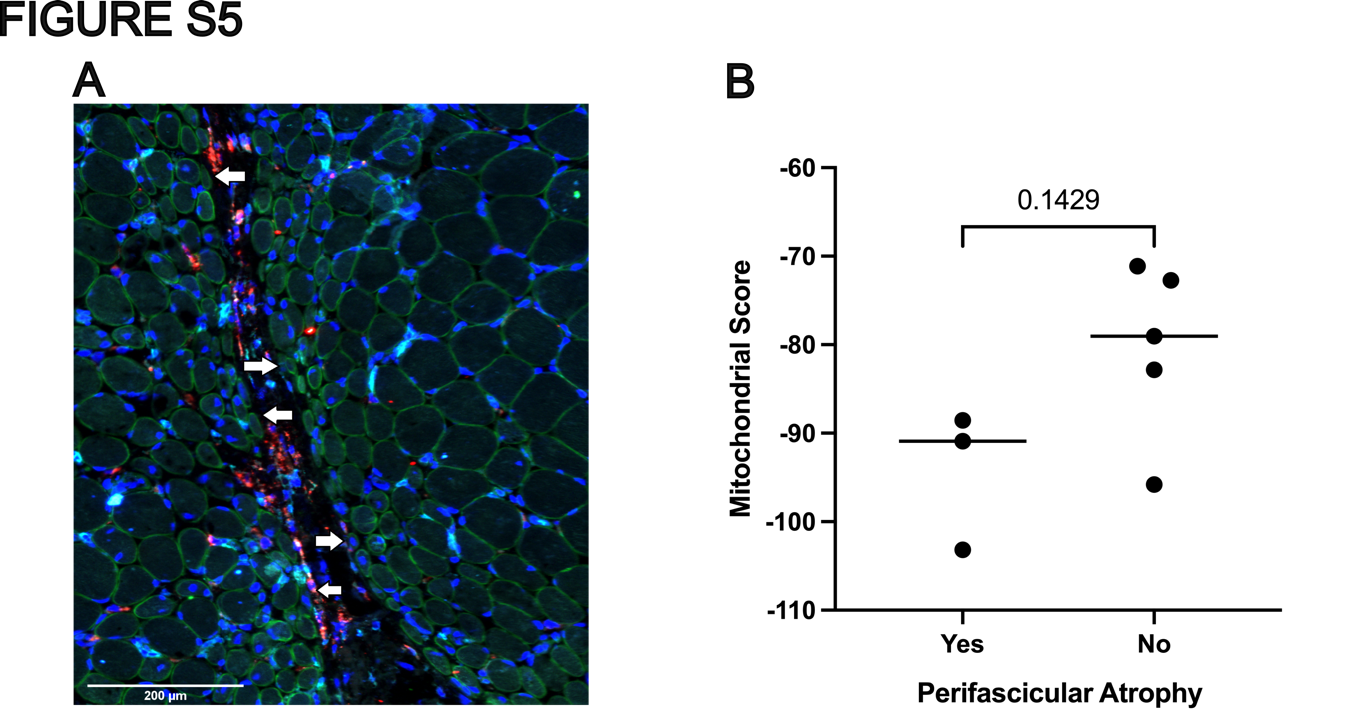

**Supplemental Figure S5. Mitochondrial gene expression in regions with and without perifascicular atrophy.** **(A)** Representative image from a JDM ROI showing a region of perifascicular atrophy. Atrophic areas were identified based on characteristic histological features. **(B)** Comparison of mitochondrial gene scores between muscle ROIs with perifascicular atrophy and those without. Scores were calculated using the 41-gene mitochondrial panel and the difference between the two groups was assessed using the Mann–Whitney U test.

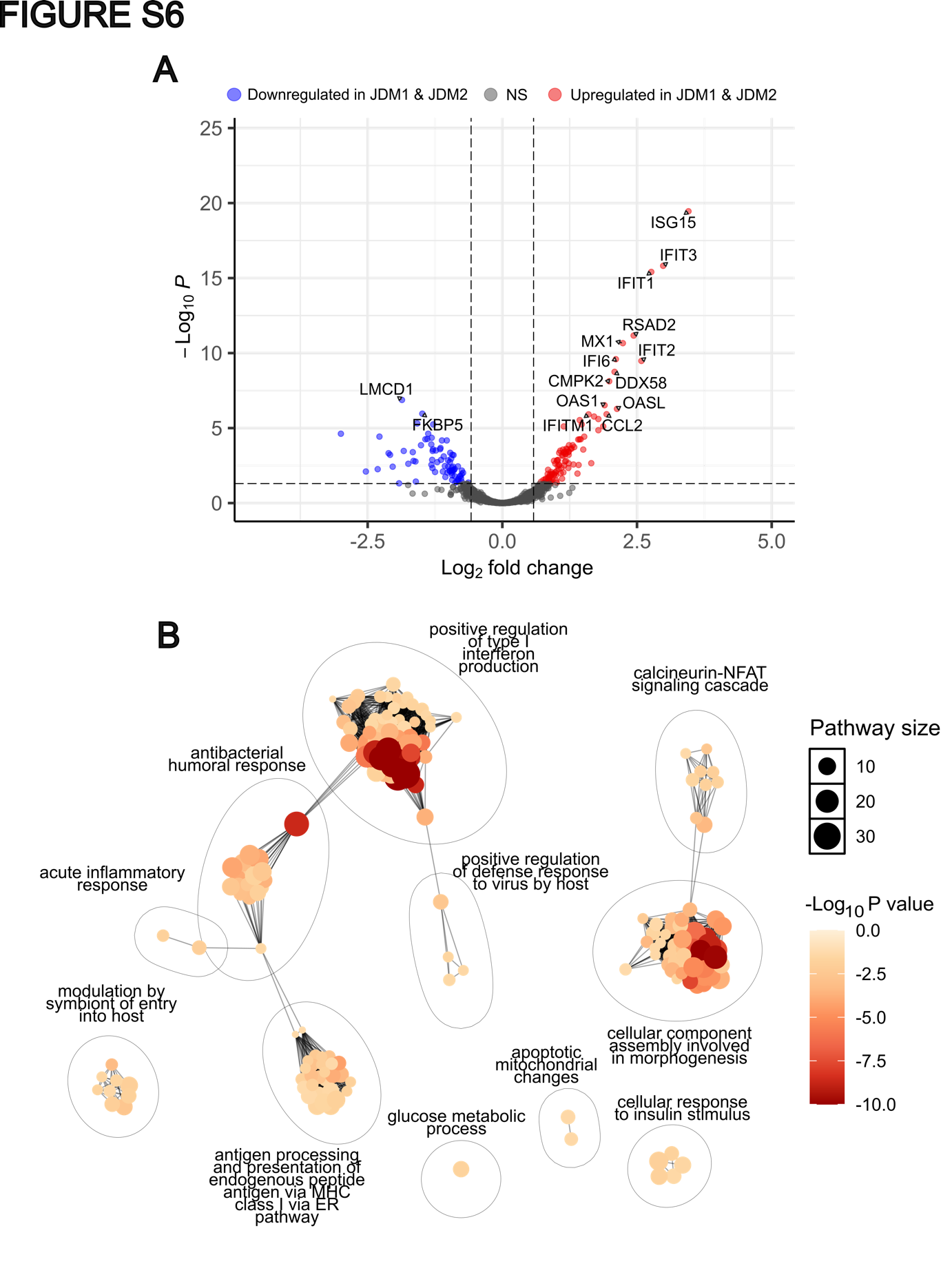

**Supplemental** **Figure S6**. **Differential gene expression analysis analysed in patients compared by muscle strength in JDM muscle ROIs.** (A) Volcano plot illustrating differentially expressed genes between patients JDM1 and JDM2 against patient JDM3 in JDM muscle ROIs. Genes (176) with an adjusted p-value ≤ 0.05 and |log2FC| ≥ 0.58 are considered statistically significant. Red points represent the upregulated genes in JDM1 and JDM2, blue points represent those downregulated. The 15 genes with the smallest adjusted p-values were annotated. (B) Network cluster plot of enriched biological pathways for the JDM1 and JDM2 versus JDM3 muscle ROI comparison, generated using the aPEAR R package. Clusters represent functionally similar pathways, based on Gene Ontology Biological Process terms, with pathways selected for inclusion based on an adjusted p-value ≤ 0.05. Larger dots represent pathways associated with a higher number of significant genes (shown in key, pathway size). Dot colour intensity corresponds to significance, with deeper red shades indicating smaller p-values (shown in colour bar, adjusted p value). NS = not significant; FC = fold change.

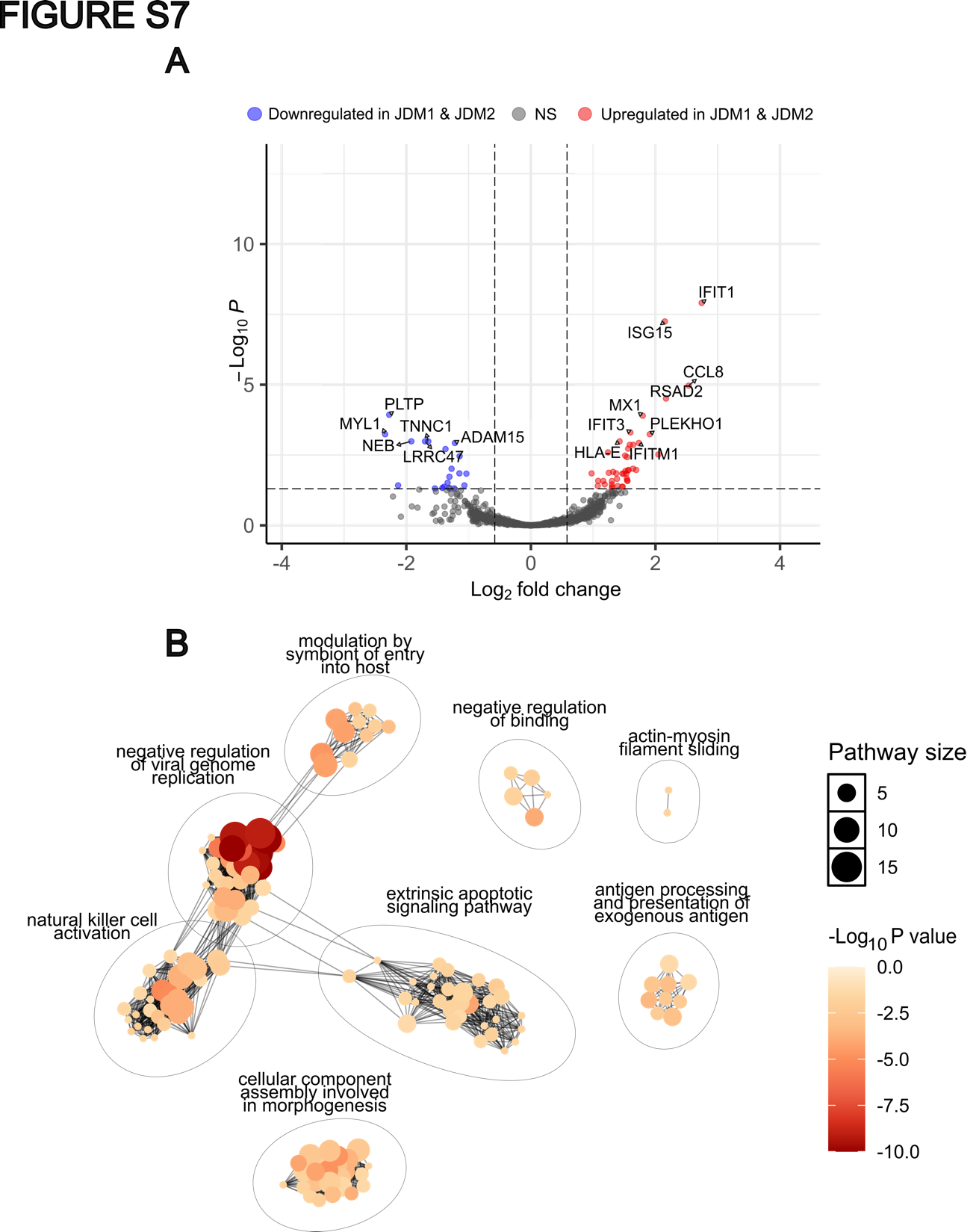

**Supplemental** **Figure S7**. **Differential gene expression analysis by muscle strength in CD68+ ROIs.** (A) Volcano plot illustrating differentially expressed genes between patients JDM1 and JDM2 against patient JDM3 in JDM CD68+ ROIs. Genes (63) with an adjusted p-value ≤ 0.05 and |log2FC| ≥ 0.58 are considered statistically significant. Red points represent the upregulated genes in JDM1 and JDM2, blue points represent those downregulated. The 15 genes with the smallest adjusted p-values were annotated. (B) Network cluster plot of enriched biological pathways for the JDM1 and JDM2 versus JDM3 CD68+ ROI comparison, generated using the aPEAR R package. Clusters represent functionally similar pathways, based on Gene Ontology Biological Process terms, with pathways selected for inclusion based on an adjusted p-value ≤ 0.05. Larger dots represent pathways associated with a higher number of significant genes (shown in key, pathway size). Dot colour intensity corresponds to significance, with deeper red shades indicating smaller p-values (shown in colour bar, adjusted p value). NS = not significant; FC = fold change.

**SUPPLEMENTAL TABLES**

**Tables S3, S5, S6, S8 and S9 are provided as a supplemental Excel document: “Supplemental_Tables_S3_S5_S6_S8_S9.xlsx”**

**Supplemental Table S1. Demographic and clinical characteristics of the JDM cohort used for histological analysis (n = 19)**

| **Characteristic** | **Value** |
| --- | --- |
| Age at Sample (years), median [IQR] | 7.24 [4.18 – 10.25] |
| Sex, n (%): |  |
| Female | 13 (62.5%) |
| Male | 6 (37.5%) |
| Disease duration at time of biopsy (months) median [IQR] | 2.46 [1.21 – 4.66] |
| Myositis specific antibody, n (%): |  |
| Nil | 4 (18.75%) |
| TIF1g | 4 (12.5%) |
| NXP2 | 4 (25%) |
| MDA5 | 4 (25%) |
| SAE | 1 (6.25%) |
| OJ | 1 (6.25%) |
| Mi2 | 1 (6.25%) |
| Disease activity at time of biopsy: |  |
| Physician global assessment score, PGA (0.0-10.0 cm), median [IQR] | 4 [2.5 – 5.8] |
| CMAS (0-52), median [IQR]* | 40.5 [11.5 – 45.25] |
| MMT8 (0-80), median [IQR]** | 48 [40 – 61] |
| CK, U/L, median [IQR] | 172 [98.5 – 1008] |
| sDAS, median [IQR] | 5 [4 – 5] |
| Muscle biopsy score data: |  |
| Inflammatory domain (0-12), median [IQR] | 4 [3 – 6] |
| Muscle fibre domain (0-10), median [IQR] | 3 [2 – 7] |
| Vascular domain (0-3), median [IQR] | 1 [0 – 1] |
| Connective tissue domain (0-2), median [IQR] | 0 [0 – 1] |
| Total Biopsy Score (0-27), median [IQR] | 8 [4.5 – 16] |
| Histopathology visual analogue score (0.0-10.0), median [IQR] | 3 [1 – 7] |

***CMAS available in n=16; was not assessed in 2 patients who were under 4 years, and was missing in 1 patient**

****MMT8 available in n=17; was not assessed in 1 patient who was under 4 years, and was missing in 1 patient**

**Supplemental Table S2. Number of ROIs sequenced per muscle section.**

| Sample ID | Healthy Muscle ROIs | JDM Muscle ROIs | JDM Muscle & CD45+ ROIs | CD68+ ROIs |
| --- | --- | --- | --- | --- |
| JDM1 |  | 3 | 1 | 2 |
| JDM2 |  | 2 | 2 | 3 |
| JDM3 |  | 2 | 2 | 5 |
| HC1 | 2 |  |  |  |
| HC2 | 2 |  |  |  |
| HC3 | 2 |  |  |  |

**Supplemental Table S3. Gene Ontology Biological Processes pathways clustered under "T cell mediated immunity" in JDM against control muscle ROIs. See: “Supplemental_Tables_S3_S5_S6_S8_S9.xlsx”**

**Supplemental Table S4. Top 20 most frequent genes recurrent across the 107 pathways clustered under “T cell mediated immunity”.**

| **Gene name** | **Number of pathways gene appears in (out of 107)** | **Comment: protein function and tissue distribution** |
| --- | --- | --- |
| *HLA-A* | 88 | MHC Class I |
| *HLA-E* | 80 | MHC Class I |
| *HLA-DRB1* | 77 | MHC Class II |
| *B2M* | 60 | MHC Class I |
| *CD81* | 57 | Tetraspanin-family member expressed on T, B cells, and in some circumstances myotubes^1^ |
| *HLA-DRA* | 54 | MHC Class II |
| *HLA-F* | 52 | MHC Class I |
| *HLA-B* | 47 | MHC Class I |
| *CD74* | 46 | HLA class II invariant chain (Ii) facilitates class ll peptide loading and transport in the ER |
| *TFRC* | 42 | Transferrin R, also known as Cluster of Differentiation 71 (CD71) important role in muscle development^2^ |
| *HLA-DPB1* | 38 | MHC Class II |
| *HLA-DPA1* | 36 | MHC Class II |
| *TAP2* | 36 | Critical for folding MHC class I, delivers cytosolic peptides to ER, binds nascent MHC class I molecules |
| *HLA-C* | 35 | MHC Class I |
| *EBI3* | 32 | Subunit of IL35 and IL27 |
| *LYN* | 29 | An SRC kinase, expressed in many tissues including muscle; upregulated in muscle in oxidative stress^3^ |
| *IL6ST* | 28 | Interleukin 6 Cytokine Family Signal Transducer, part of the IL-6 receptor, also expressed on muscle |
| *BRD2* | 23 | Nuclear serine/threonine kinase involved in transcriptional regulation also expressed in muscle |
| *NCK1* | 22 | SRC-family signalling protein involved in metabolic glucose pathways and unfolded protein response |
| *RSAD2* | 22 | An ISG expressed in many cell types in presence of IFN |

**Supplemental Table S5. fGSEA and pathway clustering of JDM against control muscle ROIs using GO CC. See “Supplemental_Tables_S3_S5_S6_S8_S9.xlsx”**

**Supplemental Table S6. fGSEA and pathway clustering of JDM against control muscle ROIs using GO BP. See “Supplemental_Tables_S3_S5_S6_S8_S9.xlsx”**

**Supplemental Table S7. List of 41 mitochondria-related genes.** The order of genes, from left to right, corresponds to their arrangement from top to bottom in the heatmap shown in Figure 4A.

| *SCO2* | *LYRM2* | *PPM1K* | *CRAT* | *ACADVL* | *CYC1* | *TOMM7* |
| --- | --- | --- | --- | --- | --- | --- |
| *ATP5ME* | *COX6B1* | *ACO2* | *UQCRB* | *COX6A2* | *COX7A1* | *ECH1* |
| *HADHB* | *NDUFA4* | *IDH2* | *CO18A* | *ATP5PO* | *COX4I1* | *ATP5F1A* |
| *MDH2* | *ATP5F1B* | *COX6C* | *SLC25A4* | *CKMT2* | *NDUFB10* | *GOT2* |
| *NDUFV1* | *NDUFS7* | *ATP5MC3* | *SLC25A11* | *CCDC51* | *ACAT1* | *ATP5MG* |
| *UQCRHL* | *ACADM* | *NNT* | *CS* | *GPT2* | *COQ10A* |  |

**Supplemental Table S8. fGSEA and pathway clustering of JDM muscle against immune cell-enriched muscle ROIs using GO CC. See “Supplemental_Tables_S3_S5_S6_S8_S9.xlsx”**

**Supplemental Table S9. fGSEA and pathway clustering of JDM muscle against immune cell-enriched muscle ROIs using GO BP. See “Supplemental_Tables_S3_S5_S6_S8_S9.xlsx”**

**Supplemental Table S10. List of antibodies and reagents used for staining GeoMx cases**

| **Name** | **Channel** | **Host** | **Company** | **Clone #** | **Catalog #** | **Dilution factor used** |
| --- | --- | --- | --- | --- | --- | --- |
| Syto13 nucleic acid stain | 488 | NA | Fisher | NA |  | 10 |
| Laminin | 532 | rabbit | Novus | Polyclonal | NB300-144AF532 | 100 |
| CD68 | 594 | mouse | Santa Cruz | KP1 | sc-20060AF594 | 400 |
| CD45 | 647 | rabbit | CST | D9M8I | 13917BF | 100 |

**METHODS**

**Patient recruitment and biopsy collection**

Patients meeting Bohan and Peter criteria for probable or definite JDM were included through the UK JDM Cohort and Biomarker Study (JDCBS)^4 5^. The study has full ethical approval through North-East Yorkshire Research Ethics Committee (MREC 01/3/022). At the time of writing 697 cases have been recruited to JDCBS of which 232 have had a diagnostic biopsy taken prior to treatment. Patient muscle biopsies (n=3) taken at diagnosis prior to treatment were selected from the JDCBS tissue biobank of biopsy samples for GeoMx analysis, based on clear infiltration of muscle tissue by CD68-positive cells^6^. In addition to the GeoMx cohort, a further 16 JDM biopsies were selected for immunohistochemical and histochemical staining for validation of key findings. Age matched control muscle tissue was obtained through the BRAIN UK Biobank with full ethical approval through South Central Hampshire Research Ethics Committee (MRE 19/SC/0217). Informed parental consent (and age-appropriate assent for children able to give assent) was obtained in accordance with the Declaration of Helsinki.

Clinical data extracted from the JDCBS Database were: demographic data, disease activity at time of biopsy, including Childhood Myositis Assessment Scale (CMAS; range 0–52; high scores indicate no weakness)^7^, manual muscle testing of 8 muscles (MMT8, range 0-80, high scores indicate no weakness)^8^, physicians global assessment (PGA range 0.0-10.0, high value indicates worse disease)^9^. Laboratory variables included myositis specific autoantibody (MSA) status, with serum tested for antibodies to TIF1g, NXP-2, MDA-5, Mi-2, SRP, HMGCR, SAE, Jo1, PL7, PL12, OJ, EJ, Zo, KS and Ha, as well as serum creatine kinase (CK), at time of biopsy.

**Histopathology**

Quadriceps muscle biopsy was obtained by open biopsy at diagnosis. Tissue was frozen and sectioned with 7μm sections. Frozen sections were stained with Haematoxylin & Eosin. Immunohistochemical staining for macrophages and T cells was performed using primary antibodies anti-human CD68 (clone PGM1) anti-human CD3 (clone UCHT1) anti-human CD20 (clone L26) as described^10^. Muscle biopsy scoring as part of routine clinical care, using the JDM Biopsy score tool was applied with assessment of the four domains and a histopathologists overall score for severity (0.0-10.0) as described^6 10^. For control cases one expert histopathologist (AM) confirmed no significant pathology in muscle tissue, including no inflammation, perifascicular atrophy or fibre necrosis.

Assessment of mitochondrial abnormalities for all 19 JDM cases was carried out using sections stained for SDH/COX as described^11^, and sections stained using primary antibodies anti-human NDUFB8 (clone 20E9DH10C12) for complex I, and anti-human MTCO1 (1clone D6E1A8) for complex IV using standard immunohistochemical staining. Biopsies were scored by one expert histopathologist (AM) who was blinded to clinical data and case details. Mitochondrial abnormality was scored as none (0), mild (1) or severe (2), based upon all three stains for assessement. For assessment of interferon-driven protein expression on muscle fibres, the same cases were stained for expression of myxovirus-resistance protein A (MxA) using primary antibody (clone M143), using the scoring system we previously published on 103 cases, scoring 0 to 3 as follows: 0 = no MxA staining; 1 = weak; 2 = moderate; 3 = strong staining, as described^12^.

**GeoMx data processing and quality control**

Tissue sections were cut from frozen samples at 7μm onto Superfrost Plus slides and formalin fixed before spatial transcriptomic analysis. The six tissue sections were analysed on the Nanostring GeoMx® DSP platform and stained for DNA and for the morphology markers laminin, CD45 and CD68 to identify muscle cells, immune cell infiltrates and tissue-resident macrophages respectively. The list of antibodies used in preparation of GeoMx analysis can be found in the online supplemental Table S10. Three types of ROIs were selected: muscle cells (from both JDM and control tissues), muscle cells with immune cell infiltrates (from JDM tissue), and regions rich in CD68-positive cells (JDM tissue). In total, 22 ROIs were obtained from JDM tissues and 6 from control tissues (see Table S2). Within the 10 ROIs focusing on infiltrating CD68+ cells, segmentation was used to focus RNA expression analysis on CD68+ cells. Using this method, the transcriptional data form CD68+ segmented ROIs will be predominantly from CD68+ cells. Defined areas of illumination (AOIs) within each CD68-enriched ROI were selected using the morphology marker CD68 and UV illumination of the AOI as described^13^.

The GeoMx Human Whole Transcriptome Atlas panel was selected, which measures 18,676 protein-encoding genes^14^, and the oligonucleotide probes were subsequently sequenced using the Illumina NovaSeq 6000, following the manufacturer's guidelines. The raw FASTQ files were processed using Nanostring’s GeoMx NGS Pipeline^15^ (version 2.3.4). To ensure data integrity, quality control to remove low-quality ROIs and negative control probes was performed using the Nanostring DSP analysis suite according to the established Nanostring pipeline. The “probeQC” counts were then extracted from the DSP suite. The R package standR^16^ was used to leverage the SpatialExperiment data infrastructure^17^ for downstream analysis of the probeQC data. Gene filtering was performed using the addPerROIQC function from standR, which removed low-expression genes while accounting for both library size and minimum count per gene. Principal component analysis (PCA) was subsequently applied for dimensionality reduction to summarize variation in the data, while relative log expression (RLE) plots were used to assess the effectiveness of data normalization.

**Differential gene expression**

Differential gene expression (DGE) analysis was conducted using the limma-voom pipeline^18^. The raw counts of the ROIs were first filtered to remove genes with low expression using the filterByExpr function from the R package edgeR^19^, thereby reducing the burden of multiple testing. Following this, the data was normalized using the voom function to account for differences in library sizes. To address potential correlations between ROIs within the same muscle section, the duplicateCorrelation function was applied. Genes were considered differentially expressed (DEGs) if they met the thresholds of |Log2 Fold Change (FC)| ≥ 0.58 (equivalent to a 1.5-fold change, a commonly used FC cutoff that balances biological significance and statistical robustness while minimizing false positives^20^) and adjusted p-value ≤ 0.05.

**Over-representation analysis**

Over-Representation Analysis (ORA) was conducted to identify biological pathways enriched among the DEGs, providing an overview of pathway dysregulation. Pathway annotations were performed using the Gene Ontology Biological Processes (GO BP)^21^ gene sets to evaluate interferon-related processes, while GO Cellular Component (GO CC)^21^ terms were used to identify mitochondrial involvement, both in conjunction with the clusterProfiler^22^ package. Pathways with an adjusted p-value ≤ 0.05 were considered statistically significant.

**Gene set enrichment analysis**

Gene set enrichment analysis (GSEA) was performed using the fgsea^23^ method, implemented with the clusterProfiler^22^ R package, to identify significantly enriched biological pathways. The gene lists for each comparison were pre-ranked based on the adjusted p-value and fold changs of each gene derived from the DGE analysis. Pathway annotation was done using the GO BP and GO CC databases^21^. Pathways with an adjusted p-value ≤ 0.05 were considered statistically significant.

**Pathway cluster network**

ORA and GSEA results were further analyzed and visualized using the R package aPEAR^24^ (Advanced Pathway Enrichment Analysis Representation). This package aids in the interpretation of pathway analysis by calculating the pairwise similarity between GO pathways, detecting clusters of overlapping pathways and assigning a biologically meaningful name for each cluster. For the ORA results, the name of each cluster was assigned based on the GO pathway with the lowest p-value, and for the GSEA results the name was assigned based on the GO pathway with the highest absolute normalized enrichment score (NES).

**Evaluation of interferon dysregulation**

To assess interferon pathway dysregulation in JDM tissues, a previously published interferon signature composed of 15 validated interferon stimulated genes (ISGs) was utilized (*LY6E, MX1, USP18, RSAD2, OAS1, IFI44L, IFI44, IFI27, ISG15, IFIT1M, IFI6, SIGLEC1, IFIT3, IRF7, STAT1*)^25^. According to the Interferome database^26^, all 15 genes are associated with both Type I and Type II interferon signalling. An interferon gene score was calculated for each ROI by summing the z-scores of the 15 interferon genes. The z-scores were computed relative to the mean and standard deviation across all ROIs, rather than controls alone, to enable comparison across distinct tissue compartments including CD68+ ROIs, which lacked control samples.

**Evaluation of mitochondrial dysfunction**

To assess mitochondrial dysfunction in JDM tissues, genes annotated with the Gene Ontology (GO) term "mitochondrion" (GO:0005739)^21^ were extracted from the Ensembl database^27^. These genes were then compared with the differentially expressed genes to identify any significant changes in mitochondrial-related gene expression. From the resulting 75 genes, 17 genes which are known ISG or can be related to IFN production were excluded including 5 genes (*IFI27, IFIT3, OAS1, RSAD2, IFI6*), which overlapped with the 15-gene interferon score. An additional 17 genes were excluded based on biological-evidence driven curation, as they were not specifically involved in mitochondrial function, (including oxidative phosphorylation, respiratory chain activity and others), mitochondrial metabolism, or mitochondrial structure. A mitochondrial gene score was calculated for each ROI by summing the z-scores of the remaining 41 genes which were tagged with the GO “mitochondrion” term. As with the interferon score, z-scores were computed relative to the mean and standard deviation across all ROIs to enable comparison across distinct tissue compartments, including CD68+ ROIs.

**Validation using published bulk RNA-sequencing data**

To independently validate findings from the spatial transcriptomic analysis, a previously published bulk RNA-sequencing dataset comprising 4 JDM muscle biopsies and 5 healthy controls was used^28^. Differential gene expression results from this dataset were obtained from the authors' published supplemental table 15 (“ST15 Muscle DEG”). From this table, significant genes overlapping with the interferon and mitochondrial gene scores identified by the GeoMx analysis were extracted for comparison. In addition, the 37 mitochondrial-encoded genes were interrogated in this validation dataset to complement findings based on the nuclear-encoded mitochondrial gene panel. GSEA was also performed using the gene list from the published results, and enrichment results were visualised using aPEAR network clustering based on combined GO Biological Process and Cellular Component annotations. Genes were considered significant if they met the thresholds of adjusted p-value ≤ 0.05 and |Log2FC| ≥ 0.58.

**Relationship between MxA and mitochondrial deficiency**

To evaluate the relationship between interferon-driven gene expression and mitochondrial dysfunction, MxA and mitochondrial deficiency scores were analysed in 19 JDM muscle biopsies (including 16 new cases and the 3 previously analysed by GeoMx). As both variables represent ordinal data, association between the two was tested using the U-statistic permutation test^29^ from the USP R package, which offers greater robustness than standard contingency table-based tests for non-parametric ordinal data. To explore associations with clinical severity, MxA and mitochondrial deficiency scores were also treated as numeric variables and correlated with clinical disease activity measurements using Spearman’s rank correlation and “BH” adjustments for multiple comparisons.

**Correlation of mitochondrial gene score with pathway-level signatures**

To determine the biological pathways most closely associated with mitochondrial dysfunction in JDM muscle, two complementary analyses were performed. First, pathway clusters from the gene set enrichment analysis comparing JDM muscle ROIs to controls were examined. For each cluster, genes that appeared as leading-edge genes in ≥75% of the pathways were selected as representative of the cluster’s core transcriptional signature. For each ROI, a cluster-specific expression score was then calculated as the sum of z-scores across the cluster's selected genes, where z-scores were computed relative to the mean and standard deviation of the gene expressions in control muscle. Scores then were computed for both muscle-only and muscle + immune-infiltrated ROIs to increase statistical power.

Second, gene set enrichment analysis was repeated for JDM muscle versus control muscle using the MSigDB Hallmark gene sets^30^. For each significantly enriched Hallmark pathway (adjusted p-value ≤ 0.05), all leading edge genes were extracted and used to calculate a Hallmark pathway score per ROI using the same z-score-based formula as above.

In both approaches, a recalculated mitochondrial gene score was generated using the same z-score method for the 41 curated mitochondrial genes, referencing the control muscle mean and standard deviation. Pearson’s correlations were then performed between the mitochondrial score and each cluster-specific or Hallmark pathway score, as appropriate. Multiple testing correction was applied using the Benjamini-Hochberg (BH) method separately for each analytical approach.

**Analysis of variation between groups**

To assess variation across groups, data were first assessed for normality using the Shapiro–Wilk test (p > 0.05 indicating a normal distribution). For comparisons involving normally distributed data, one-way ANOVA was used, followed by Tukey’s post-hoc test for pairwise comparisons, which includes correction for multiple testing. For non-normally distributed data (Shapiro-Wilk p ≤ 0.05), the Kruskal–Wallis test was applied, followed by Dunn’s post-hoc test with adjustments for multiple comparisons.

**Data analysis**

All statistical modelling, gene expression, pathway and network analyses were conducted using R (version 4.4.1, https://github.com/WedderburnLab/GeoMx-pipeline) and RStudio (version 2024.12.0+467), with data visualizations and additional statistical comparisons produced using both R and GraphPad Prism (version 10.3.1).

**Acknowledgements**

The Juvenile Dermatomyositis Cohort Biomarker Study & Repository (JDCBS) would like to thank all of the patients and their families who contributed to the JDCBS research study. We thank all local research coordinators and principal investigators who have made this research possible. Clinical, research and administrative contributors to JDCBS members were as follows:

Dr Kate Armon, and Ms Louise Coke, Ms Julie Cook and Ms Amy Nichols (Norfolk and Norwich University Hospitals);Dr Liza McCann, Mr Ian Roberts, Dr Eileen Baildam, Ms Louise Hanna, Ms Olivia Lloyd, Susan Wadeson, Ms Michelle Andrews, Ms Olivia Lloyd, Mrs Jane Roach and Dr Beverley Almeida (The Royal Liverpool Children’s Hospital, Alder Hey, Liverpool); Dr Phil Riley, Ms Ann McGovern, and Ms Verna Cuthbert (Royal Manchester Children’s Hospital, Manchester); Dr Clive Ryder, Ms Janis Scott, Ms Beverley Thomas, Professor Taunton Southwood, Dr Eslam Al-Abadi and Ms Ruth Howman (Birmingham Children’s Hospital, Birmingham); Dr Sue Wyatt, Mrs Gillian Jackson, Dr Mark Wood, Dr Tania Amin, Dr Vanessa VanRooyen, Ms Deborah Burton, Ms Louise Turner, Ms Heather Rostron, and Ms Sarah Hanson (Leeds General Infirmary, Leeds); Dr Joyce Davidson, Dr Janet Gardner-Medwin, Dr Neil Martin, Ms Sue Ferguson, Ms Liz Waxman and Mr Michael Browne, Ms Roisin Boyle, Ms Emily Blyth, Ms Susanne Cathcart, Dr Kirsty McLellan and Dr Jaclyn Keightley (The Royal Hospital for Sick Children, Yorkhill, Glasgow); Dr Mark Friswell, Professor Helen Foster, Ms Alison Swift, Dr Sharmila Jandial, Ms Vicky Stevenson, Ms Debbie Wade, Dr Ethan Sen, Dr Eve Smith, Ms Lisa Qiao, Mr Stuart Watson and Ms Claire Duong, Dr Stephen Crulley, Mr Andrew Davies, Miss Caroline Miller, Ms Lynne Bell, Dr Flora McErlane, Dr Sunil Sampath, Dr Josh Bennet, Mrs Sharon King, Mr Christopher Long and Ms Lesley Brindley (Great North Children’s Hospital, Newcastle); Dr Helen Venning, Dr Rangaraj Satyapal, Mrs Elizabeth Stretton, Ms Mary Jordan, Dr Ellen Mosley, Ms Anna Frost, Ms Lindsay Crate, Dr Kishore Warrier, Ms Stefanie Stafford, Mrs Brogan Wrest, Ms Chia-Ping Chou, and Mr Paul Pryce (Queens Medical Centre, Nottingham); Professor Lucy Wedderburn, Dr Clarissa Pilkington, Dr Nathan Hasson, Dr Muthana Al-Obadi, Dr Giulia Varnier, Dr Sandrine Lacassagne, Ms Sue Maillard, Mrs Lauren Stone, Ms Elizabeth Halkon, Ms Virginia Brown, Ms Audrey Juggins, Dr Sally Smith, Ms Sian Lunt, Ms Elli Enayat, Ms Hemlata Varsani, Ms Laura Kassoumeri, Miss Laura Beard, Ms Katie Arnold, Mrs Yvonne Glackin, Ms Stephanie Simou, Dr Beverley Almeida, Dr Kiran Nistala, Dr Raquel Marques, Dr Claire Deakin, Dr Parichat Khaosut, Ms Stefanie Dowle, Dr Charalampia Papadopoulou, Dr Shireena Yasin, Dr Christina Boros, Dr Meredyth Wilkinson, Dr Chris Piper, Ms Cerise Johnson-Moore, Ms Lucy Marshall, Ms Kathryn O’Brien, Ms Emily Robinson, Mr Dominic Igbelina, Dr Polly Livermore, Dr Socrates Varakliotis, Ms Rosie Hamilton, Ms Lucy Nguyen, Mr Dario Cancemi, Dr Ovgu Kul Cinar, Dr Elena Moraitis, Dr Hannah Peckham and Dr Qiong Wu (Great Ormond Street Hospital, London); Dr Kevin Murray (Princess Margaret Hospital, Perth, Western Australia); Dr Coziana Ciurtin, Dr John Ioannou, Mrs Caitlin Clifford, Ms Linda Suffield and Ms Laura Hennelly (University College London Hospital, London); Ms Helen Lee, Ms Sam Leach, Ms Helen Smith, Dr Anne-Marie McMahon, Ms Heather Chisem, Ms Jeanette Hall and Ms Amy Huffenberger (Sheffield’s Children’s Hospital, Sheffield); Dr Nick Wilkinson, Ms Emma Inness, Ms Eunice Kendall, Mr David Mayers, Ms Ruth Etherton, Ms Danielle Miller and Dr Kathryn Bailey (Oxford University Hospitals, Oxford); Dr Jacqui Clinch, Ms Natalie Fineman, Ms Helen Pluess-Hall, Ms Suzanne Sketchley, Ms Melanie Marsh, Ms Anna Fry, Ms Maisy Dawkins-Lloyd and Ms Mashal Asif (Bristol Royal Hospital for Children, Bristol); Dr Joyce Davidson, Margaret Connon and Ms Lindsay Vallance (Royal Aberdeen Children’s Hospital); Dr Kirsty Haslam, Ms Charlene Bass-Woodcock, Ms Trudy Booth, and Ms Louise Akeroyd (Bradford Teaching Hospitals); Dr Alice Leahy, Amy Collier, Rebecca Cutts, Emma Macleod, Dr Hans De Graaf, Dr Brian Davidson, Sarah Hartfree, Ms Elizabeth Fofana and Ms Lorena Caruana (University Hospital Southampton); Dr Catriona Anderson (Royal Hospital for Children and Young People, Edinburgh); Dr Jayne MacMahon and Dr Peter Bale (Cambridge University Hospitals); and all the Children, Young people and their families who have contributed to this research.

**SUPPLEMENTARY REFERENCES**

1. Charrin S, Latil M, Soave S, et al. Normal muscle regeneration requires tight control of muscle cell fusion by tetraspanins CD9 and CD81. *Nat Commun* 2013;4:1674. doi: 10.1038/ncomms2675

2. Li Y, Cheng JX, Yang HH, et al. Transferrin receptor 1 plays an important role in muscle development and denervation-induced muscular atrophy. *Neural Regen Res* 2021;16(7):1308-16. doi: 10.4103/1673-5374.301024

3. Yang J, Zhai Y, Huang C, et al. RP105 Attenuates Ischemia/Reperfusion-Induced Oxidative Stress in the Myocardium via Activation of the Lyn/Syk/STAT3 Signaling Pathway. *Inflammation* 2024;47(4):1371-85. doi: 10.1007/s10753-024-01982-y [published Online First: 20240403]

4. Martin N, Krol P, Smith S, et al. A national registry for juvenile dermatomyositis and other paediatric idiopathic inflammatory myopathies: 10 years' experience; the Juvenile Dermatomyositis National (UK and Ireland) Cohort Biomarker Study and Repository for Idiopathic Inflammatory Myopathies. *Rheumatology* 2010;50(1):137-45. doi: 10.1093/rheumatology/keq261

5. <https://juveniledermatomyositis.org.uk/jdrg/>.

6. Varsani H, Charman SC, Li CK, et al. Validation of a score tool for measurement of histological severity in juvenile dermatomyositis and association with clinical severity of disease. *Ann Rheum Dis* 2015;74(1):204-10. doi: 10.1136/annrheumdis-2013-203396 [published Online First: 20130924]

7. Huber AM, Feldman BM, Rennebohm RM, et al. Validation and clinical significance of the Childhood Myositis Assessment Scale for assessment of muscle function in the juvenile idiopathic inflammatory myopathies. *Arthritis Rheum* 2004;50(5):1595-603. doi: 10.1002/art.20179

8. Rider LG, Koziol D, Giannini EH, et al. Validation of manual muscle testing and a subset of eight muscles for adult and juvenile idiopathic inflammatory myopathies. *Arthritis Care Res (Hoboken)* 2010;62(4):465-72. doi: 10.1002/acr.20035

9. Rider LG, Feldman BM, Perez MD, et al. Development of validated disease activity and damage indices for the juvenile idiopathic inflammatory myopathies: I. Physician, parent, and patient global assessments. Juvenile Dermatomyositis Disease Activity Collaborative Study Group. *Arthritis Rheum* 1997;40(11):1976-83. doi: 10.1002/art.1780401109

10. Wedderburn LR, Varsani H, Li CK, et al. International consensus on a proposed score system for muscle biopsy evaluation in patients with juvenile dermatomyositis: a tool for potential use in clinical trials. *Arthritis Rheum* 2007;57(7):1192-201. doi: 10.1002/art.23012

11. Dubowitz V, Sewry CA, Oldfors A. Muscle biopsy: a practical approach. Muscle biopsy: a practical approach. London: Elsevier 2020:14-23.

12. Soponkanaporn S, Deakin CT, Schutz PW, et al. Expression of myxovirus-resistance protein A: a possible marker of muscle disease activity and autoantibody specificities in juvenile dermatomyositis. *Neuropathol Appl Neurobiol* 2019;45(4):410-20. doi: 10.1111/nan.12498 [published Online First: 20180604]

13. Böning S, Schneider F, Huber AK, et al. Region of interest localization, tissue storage time, and antibody binding density-a technical note on the GeoMx® Digital Spatial Profiler. *Immunooncol Technol* 2024;23:100727. doi: 10.1016/j.iotech.2024.100727 [published Online First: 20240820]

14. Oszwald A, Zisser L, Schachner H, et al. Full-length target sequences of GeoMx digital spatial profiling probes reveal that gene-promiscuity predicts probe sensitivity to EDTA tissue decalcification. *Scientific Reports* 2024;14(1):21156. doi: 10.1038/s41598-024-72335-w

15. Merritt CR, Ong GT, Church SE, et al. Multiplex digital spatial profiling of proteins and RNA in fixed tissue. *Nat Biotechnol* 2020;38(5):586-99. doi: 10.1038/s41587-020-0472-9 [published Online First: 20200511]

16. Liu N, Bhuva DD, Mohamed A, et al. standR: spatial transcriptomic analysis for GeoMx DSP data. *Nucleic Acids Res* 2024;52(1):e2. doi: 10.1093/nar/gkad1026

17. Righelli D, Weber LM, Crowell HL, et al. SpatialExperiment: infrastructure for spatially resolved transcriptomics data in R using Bioconductor. *bioRxiv* 2022:2021.01.27.428431. doi: 10.1101/2021.01.27.428431

18. Law CW, Chen Y, Shi W, et al. voom: Precision weights unlock linear model analysis tools for RNA-seq read counts. *Genome Biol* 2014;15(2):R29. doi: 10.1186/gb-2014-15-2-r29 [published Online First: 20140203]

19. Chen Y, Chen L, Lun ATL, et al. edgeR 4.0: powerful differential analysis of sequencing data with expanded functionality and improved support for small counts and larger datasets. *bioRxiv* 2024:2024.01.21.576131. doi: 10.1101/2024.01.21.576131

20. Dalman MR, Deeter A, Nimishakavi G, et al. Fold change and p-value cutoffs significantly alter microarray interpretations. *BMC Bioinformatics* 2012;13 Suppl 2(Suppl 2):S11. doi: 10.1186/1471-2105-13-s2-s11 [published Online First: 20120313]

21. Consortium TGO. The Gene Ontology resource: enriching a GOld mine. *Nucleic Acids Research* 2020;49(D1):D325-D34. doi: 10.1093/nar/gkaa1113

22. Wu T, Hu E, Xu S, et al. clusterProfiler 4.0: A universal enrichment tool for interpreting omics data. *Innovation (Camb)* 2021;2(3):100141. doi: 10.1016/j.xinn.2021.100141 [published Online First: 20210701]

23. Korotkevich G, Sukhov V, Budin N, et al. Fast gene set enrichment analysis. *bioRxiv* 2021:060012. doi: 10.1101/060012

24. Kerseviciute I, Gordevicius J. aPEAR: an R package for autonomous visualization of pathway enrichment networks. *Bioinformatics* 2023;39(11) doi: 10.1093/bioinformatics/btad672

25. Wilkinson MGL, Moulding D, McDonnell TCR, et al. Role of CD14+ monocyte-derived oxidised mitochondrial DNA in the inflammatory interferon type 1 signature in juvenile dermatomyositis. *Ann Rheum Dis* 2023;82(5):658-69. doi: 10.1136/ard-2022-223469 [published Online First: 20221223]

26. Rusinova I, Forster S, Yu S, et al. INTERFEROME v2.0: an updated database of annotated interferon-regulated genes. *Nucleic Acids Research* 2012;41(D1):D1040-D46. doi: 10.1093/nar/gks1215

27. Harrison PW, Amode MR, Austine-Orimoloye O, et al. Ensembl 2024. *Nucleic Acids Res* 2024;52(D1):D891-d99. doi: 10.1093/nar/gkad1049

28. Roberson EDO, Mesa RA, Morgan GA, et al. Transcriptomes of peripheral blood mononuclear cells from juvenile dermatomyositis patients show elevated inflammation even when clinically inactive. *Sci Rep* 2022;12(1):275. doi: 10.1038/s41598-021-04302-8 [published Online First: 20220107]

29. Thomas BB, Ioannis K, Richard JS. Optimal rates for independence testing via <i>U</i>-statistic permutation tests. *The Annals of Statistics* 2021;49(5):2457-90. doi: 10.1214/20-AOS2041

30. Liberzon A, Birger C, Thorvaldsdóttir H, et al. The Molecular Signatures Database (MSigDB) hallmark gene set collection. *Cell Syst* 2015;1(6):417-25. doi: 10.1016/j.cels.2015.12.004
